# Implications of trade network structure and population dynamics for food security and equality

**DOI:** 10.1101/2021.07.08.451671

**Authors:** Kathyrn R Fair, Chris T Bauch, Madhur Anand

## Abstract

Given trade’s importance to maintaining food security, it is crucial to understand the relationship between human population growth, land use, food supply, and trade. We develop a metapopulation model coupling human population dynamics to agricultural land use and food production in “patches” (regions and countries) connected via trade networks. Patches that import sparingly or fail to adjust their demand sharply in response to changes in food per capita experience food insecurity. They fall into a feedback loop between increasing population growth and decreasing food per capita, particularly if they are peripheral to the network. A displacement effect is also evident; patches that are more central and/or import more heavily preserve their natural land states. Their reliance on imports means other patches must expand their agricultural land. These results emphasize that strategies for improving food security and equality must account for the combined effects of network topology and patch-level characteristics.

---

The global population is expected to grow to 10.9 billion by 2100 [1]. Producing enough food for this population, given predicted shifts in consumption patterns, is a challenge emphasized in the literature [2, 3, 4, 5]. An estimated 60% increase in food production by 2050 (from 2006/07) will be needed to meet growing demand [6]. This increase may be achieved partially through higher yields and agricultural intensification [2, 7], however agricultural land expansion will also be required [8, 7, 3, 9, 10, 11, 2]. The necessity of expansion raises concerns due to the environmental impact of converting natural land states (e.g. forest) for agriculture and the finite amount of land suitable for agriculture, some of which will be lost to urbanization and land degradation [2, 3, 5]. To practice sustainable development, understanding how human population growth interacts with the global food system to impact land use and food security is necessary [12, 13, 5].

A crucial aspect of the global food system is trade. Around 23% of food calories produced for human consumption move through the global trade network [14]. From 1986-2009 the amount of calories traded increased by more than 200%, with a more than 50% increase in the number of edges (links) in the network [14]. Trade impacts human population growth, food security, land use changes, and the environmental impacts of food production [2, 3, 15, 16, 14]. These effects may be positive (e.g. efficient distribution of food), or negative (e.g. increased agricultural exports leading to accelerated deforestation) [2, 15].

The coupled human-environment system created by the interplay between population growth, land use, and food trade/production can be conceptualized by representing human populations as components of a metapopulation. A metapopulation is a set of spatially distinct populations of the same species that interact through the migration of individuals between populations [17, 18]. In addition to exchanging individuals, human populations exchange resources through trade. Mathematical models can be used to explore the dynamics of human metapopulations, helping us gain insight regarding the effects of migration [19, 20], conflict and bargaining over common resources [21, 22, 23], and resource transfer [19, 24, 25, 26]. This latter aspect forms the focus of our study.

Our objective is to explore how metapopulation structure – not only the inclusion of trade, but trade network structure and the parameters controlling demand-driven trade dynamics – impacts a system where human population growth, food supply, and agricultural land area are coupled. We develop a differential equation model based on previous work exploring human metapopulations [24] and land use change [8]. Our model includes features not shared by similar models in the published literature, with respect to use of empirical data and model structure. Firstly, our model is parametrized with empirical data and fit to historical trends in global agricultural land use, food supply, and population growth. Secondly, our model allows us to explore how trade impacts population growth while accounting for agricultural land use. Thirdly, our model incorporates both resource-limited population growth in low-resource settings [24, 19, 25, 26] and the observed demographic transition towards lower fertility rates in high-resource settings (developed economies) [27, 28, 29].

## Results

Our metapopulation model couples human population dynamics to food production and agricultural land use (Fig. 1). Agricultural land in each patch (representing a region or country) produces food. Some of this food is consumed domestically, while the rest is redistributed throughout the metapopulation via a trade network. Food supply (obtained through domestic production and imports) and the current population size influence the net population growth rate. Food demand within each patch (a function of food per capita) drives both land use change (expansion of agricultural land) and food imports, determining available food. We simulate this model to explore how trade network structure and patch-level characteristics impact outcomes.

**Figure 1.**
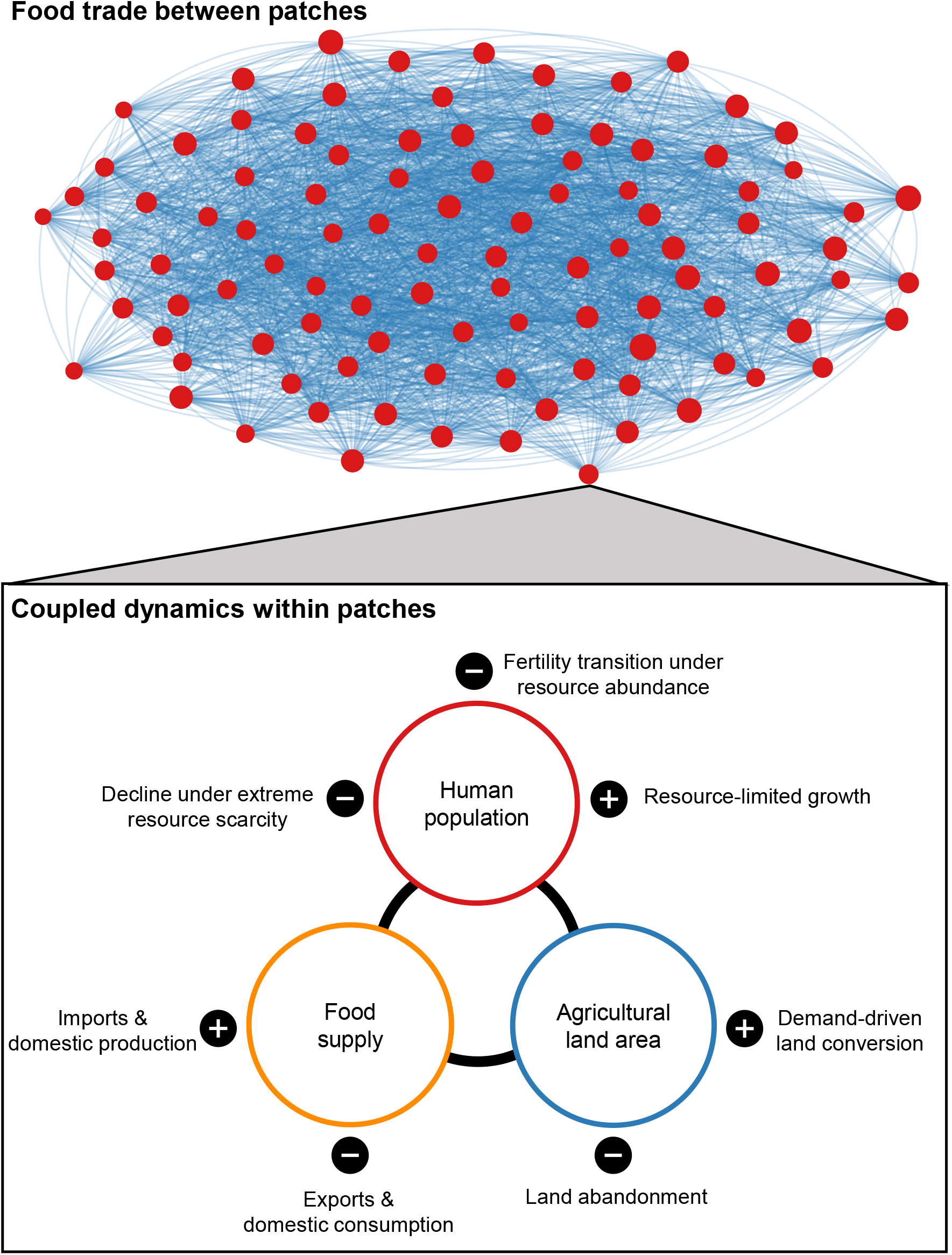
Schematic representation of metapopulation model. Mechanisms causing increases in a quantity (population, agricultural land area, or food supply) are indicated by a +, while those resulting in decreases are indicated by a –. Agricultural yield (which increases towards a maximum value over time) as well as agricultural land area determine domestic food production volumes. See Methods for model equations.

The characteristics we focus on are patch-level food demand thresholds (controlled by *β*: lower *β*-values indicate a higher threshold and thus higher demand for domestic and/or imported food, at a given level of food availability per capita) and responsiveness to changes in food per capita (controlled by *γ*: higher *γ*-values indicate a sharp change in demand for domestic or imported food, for a given change in food availability per capita). Outcomes are characterised by model variables (population size, agricultural land area, and food supply) as well as natural land state area (all land not in use for agriculture or urban area), patch level food per capita (FPC) and non-agricultural land (urban area + natural land states) per capita (LPC) and Gini index values for these per capita amounts. Gini index values range between 0 and 1, with larger values indicating higher levels of inequality.

### Demand thresholds and responsiveness

We first consider how patch-level food demand thresholds and responsiveness to changes in food per capita impact outcomes. When patches adjust their demand sharply in response to changes in food per capita – i.e. have high responsiveness (*γ*) – change within the system is accelerated. High food demand, resulting from high demand thresholds (low *β*-values), leads to between-patch inequalities that cannot be rectified by increasing responsiveness (unlike when thresholds are low).

Heightened responsiveness shifts the system from a region of parameter space where population, agricultural land, natural land state area, and food supply (Fig. 2a-f) have moderate values, to a regime where extreme outcomes are possible. Here, high responsiveness means that we observe starkly different outcomes within the time-frame considered, depending on the demand threshold (*β*-value). Higher responsiveness also reduces inequality (Fig. 2m-p) when the demand threshold is low (*β* > 1). However, if the threshold is high (*β* < 1) the resultant high levels of inequality cannot reduced by increasing responsiveness. When both responsiveness and demand thresholds are high (large *γ*, small *β*) the resulting global population is small, and has high mean food and non-agricultural land per capita (Fig. 2a,b, i-l), however these resources are unequally distributed (Fig. 2m-p).

**Figure 2.**
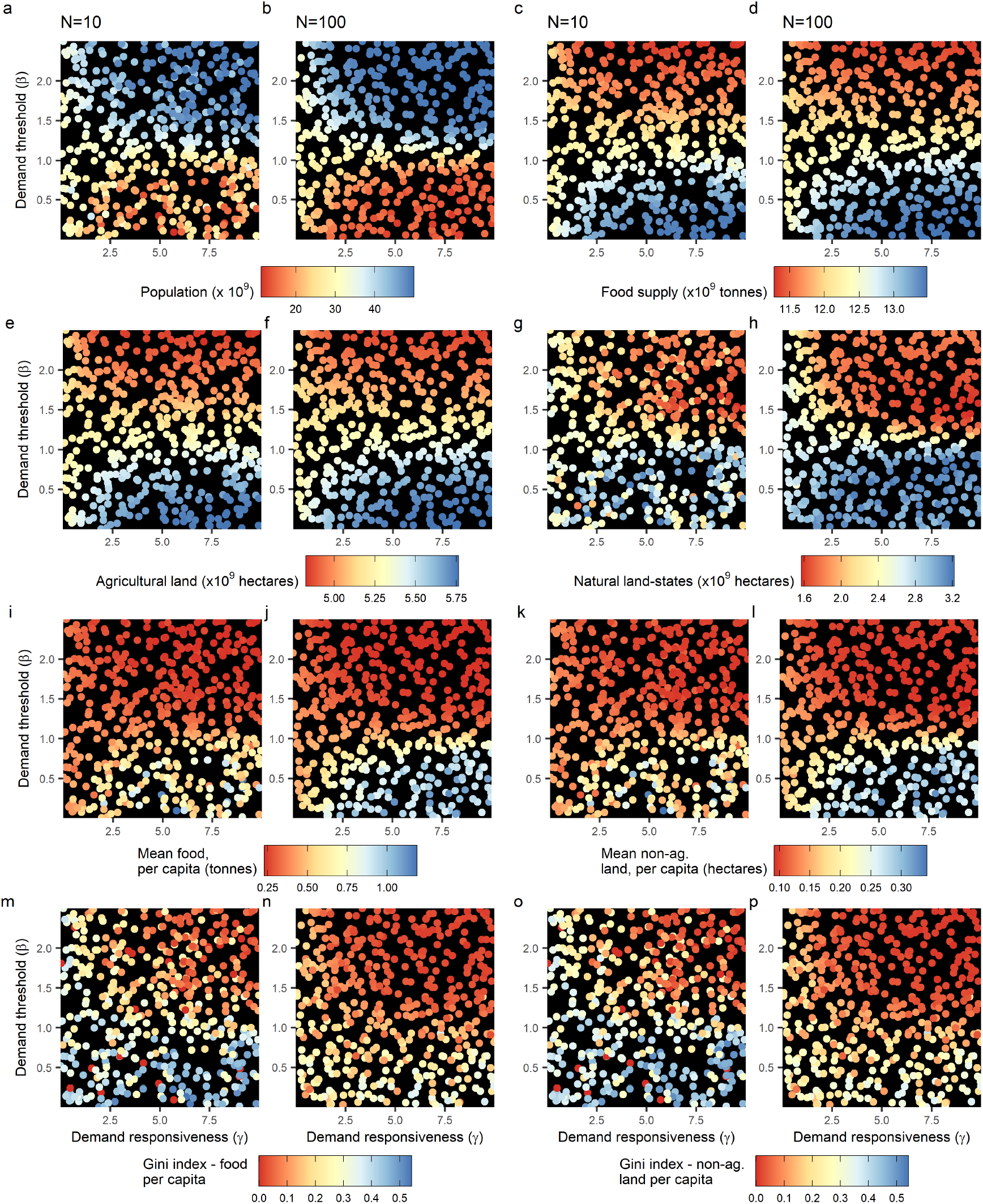
When food demand is high (*β* is small), inequality between patches will not be reduced if patches adjust their demand more sharply in response to changes in food per capita (i.e. if *γ* is increased). Figure panels show how, under a low yield scenario, demand responsiveness (*γ*) and threshold (*β*) impact global a)-b) population c)-d) food supply, e)-f) agricultural land area, g)-h) natural land state area, as well as mean patch-level per capita i)-j) food and k)-l) non-agricultural land, and Gini index, per capita m)-n) food and o)-p) non-agricultural land. Each point indicates a realization of the model. Model parameter settings (except for *β*, *γ*) appear in Supplementary information, Table S1.

Outcomes are qualitatively similar for 10 and 100 patch networks, though for larger networks (*N* = 100) demarcations between regions where different outcomes are possible are more sharply defined, while smaller (*N* = 10) networks show more variation in inequality (Fig. 2i-l). Trading on larger networks reduces inequality (for a given combination of responsiveness and threshold). Results are similar under a high yield scenario (Supplementary information, Fig. S5) though as increased yield expand the global food supply, the global population size will be lower (due to the transition to lower growth rates in food-rich populations). The remainder of our analyses focus on the low yield scenario.

### Food sources

We experiment with a scenario where demand for the 2 possible sources of food (domestically produced and imported) differs i.e. there are separate demand thresholds (*β*-values) for food from domestic production (*β_A_*, which drives agricultural land expansion) and from imports (*β^I^*). The demand threshold for domestically produced food (where demand for domestic food drives agricultural land expansion) is a strong determinant of outcomes (Fig. 3).

**Figure 3.**
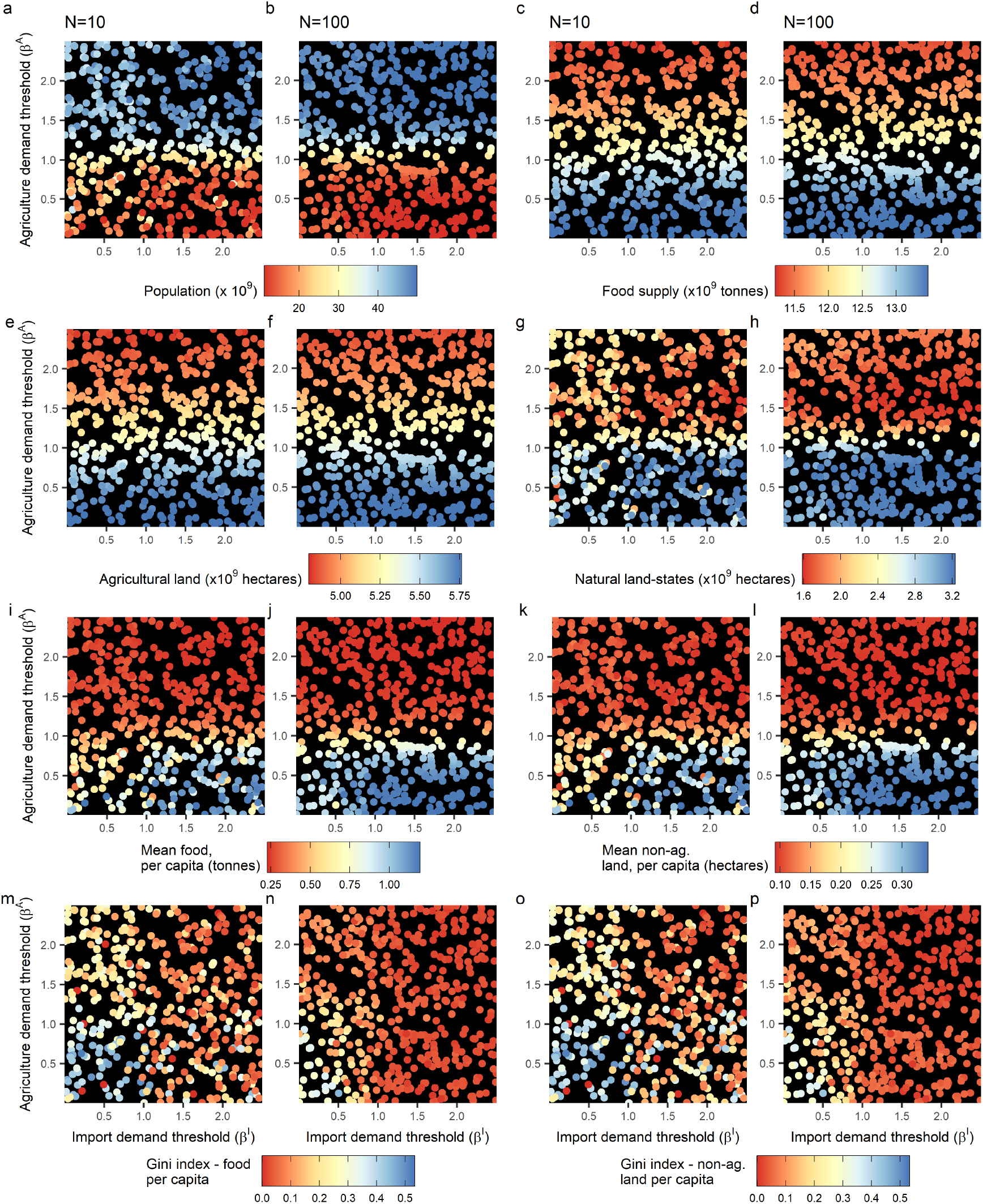
Agricultural land expansion, driven by demand for domestically produced food (higher when *β^A^* is smaller), is a strong limiting factor on global population size regardless of trade (i.e. regardless of *β^I^*). Figure panels show how the demand thresholds for different food sources impacts global a)-b) population, c)-d) food supply, e)-f) agricultural land area, g)-h) natural land state area, as well as mean patch-level per capita i)-j) food and k)-l) non-agricultural land, and Gini index, per capita m)-n) food and o)-p) non-agricultural land. Each point indicates a realization of the model. Model parameter settings (except for *γ* = 7.5, *β^A,I^*) appear in Supplementary information, Table S1.

If the demand threshold for domestically produced food is low (*β^A^* > 1), patches will only expand their agricultural land area when food per capita is very low, and the sole avenue for increased domestic food production in food-rich patches is improved yield. As yield is bounded above by some theoretical maximum, and increases at a much slower rate than land can be converted, this severely limits the rate at which food production can be increased. These limitations on food supply lessen the impact of trade on the system, as the import demand threshold (*β^I^*) only determines how food is re-distributed through the trade network. The resulting global population is large and experiences low levels of inequality, however this comes at the cost of extremely low levels of natural land states, and of food and non-agricultural land per capita. If the demand threshold for domestically produced food is high (*β^A^* < 1), food supply increases quickly due to agricultural land expansion and trade has a larger impact on system dynamics. Here, if the demand threshold for imported food is also high (*β^I^* < 1) inequality is higher and mean food and non-agricultural land per capita are lower. Inequalities resulting from high thresholds (*β^A^*, *β^I^* < 1) are more substantial in smaller networks.

### Network structure

We examine how trade network structure impacts outcomes, where trade may occur on more regularly structured networks (low values for *p*, the rewiring probability), small-world networks (intermediate *p*-values), or more randomly structured networks (high *p*-values). More random structures promote inequality, with Gini index values increasing substantially as *p* increases (Fig. 4g,h).

**Figure 4.**
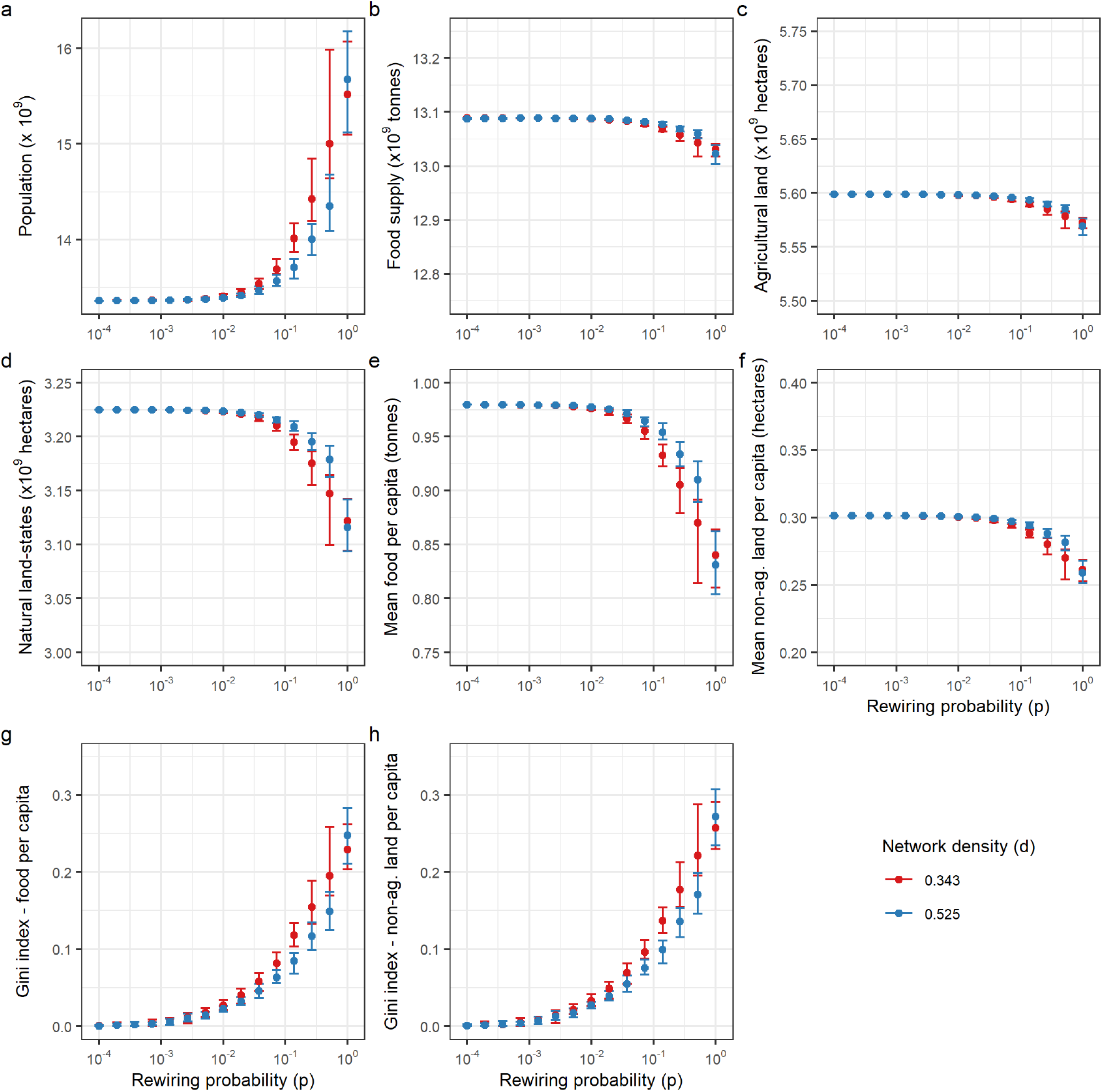
More random network structures, occurring when *p* is large, promote inequality. Figure panels show the effect of increasing the randomness of the network structure (increasing *p*) on global a) population b) food supply c) agricultural land area d) natural land state area, as well as mean per capita e) food and f) non-agricultural land, and Gini index, per capita g) food and h) non-agricultural land. Bars indicate the range of outcomes (min/max values) and points indicate mean values across all realizations of the model. Model parameter settings (except for *γ* = 7.5, *β* = 0.75) appear in Supplementary information, Table S1.

Metapopulations trading on more randomly structured networks have larger populations than those trading on more regular networks (Fig. 4a) though values for agricultural land, food supply, and mean non-agricultural land per capita are similar regardless of network structure (Fig. 4b-c,f). As a result of the relatively stable food supply and larger population sizes as *p* is increased, metapopulations trading on more random networks will experience lower levels of mean food per capita and smaller areas of natural land states (Fig. 4d,e). These results hold at both network densities, though for high *p*-values metapopulations on higher density networks may experience slightly better outcomes; i.e. lower levels of inequality, higher mean food per capita, etc. Additionally, the relative position of patches (located on nodes) within the network impacts their outcomes; patches situated more centrally within the network are able to obtain higher levels of food per capita than those located on the network’s periphery (Supplementary information, Fig. S4).

### Heterogeneity in patch-level characteristics

As it is unlikely that real-world metapopulations will be homogeneous in their characteristics, we explore the impact of differences in patch-level demand responsiveness and import demand threshold. This is done within a simplified framework where there are 2 possible levels for each characteristic (low vs. high). These experiments indicate that heterogeneity in these characteristics exacerbates inequalities resulting from network topology, and negatively impacts food security.

#### Import demand threshold

When patch-level import demand thresholds 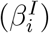 are heterogeneous (50% of patches have high thresholds, 50% have low thresholds), patches that import heavily (i.e. have high thresholds) experience better outcomes than those that import sparingly (i.e. have low thresholds), resulting in high levels of inequality. We note that the volume of food obtained via import depends not only on demand for imports, but also on the availability of food for import. Heavily and sparingly should be understood as referring to the amount of imported food obtained relative to the total amount available via import, as opposed to the absolute amount obtained.

In this heterogeneous scenario, a spike in global population occurs, accompanied by decreases in mean per capita food levels and increases in inequality (Fig. 5a,b,c), though both global food supply and agricultural land area are roughly the same regardless of the distribution of threshold values (Supplementary information, Fig. S6). Initially increasing trajectories for mean food per capita do not persist, with a pronounced downturn under the 50/50 parametrization. Inequalities in the 50/50 scenario stem from the fact that, though patches that import heavily (i.e. have high thresholds) rapidly obtain high food per capita and undergo a demographic transition, patches that import sparingly (i.e. have low thresholds) get caught in a feedback loop between decreasing food per capita and increasing population growth rates (Supplementary information, Fig. S7). This leads to marked differences in food per capita (and population size) for patches with differing import thresholds at *t* = 100 (Fig. 5d, Supplementary information, Fig. S8).

**Figure 5.**
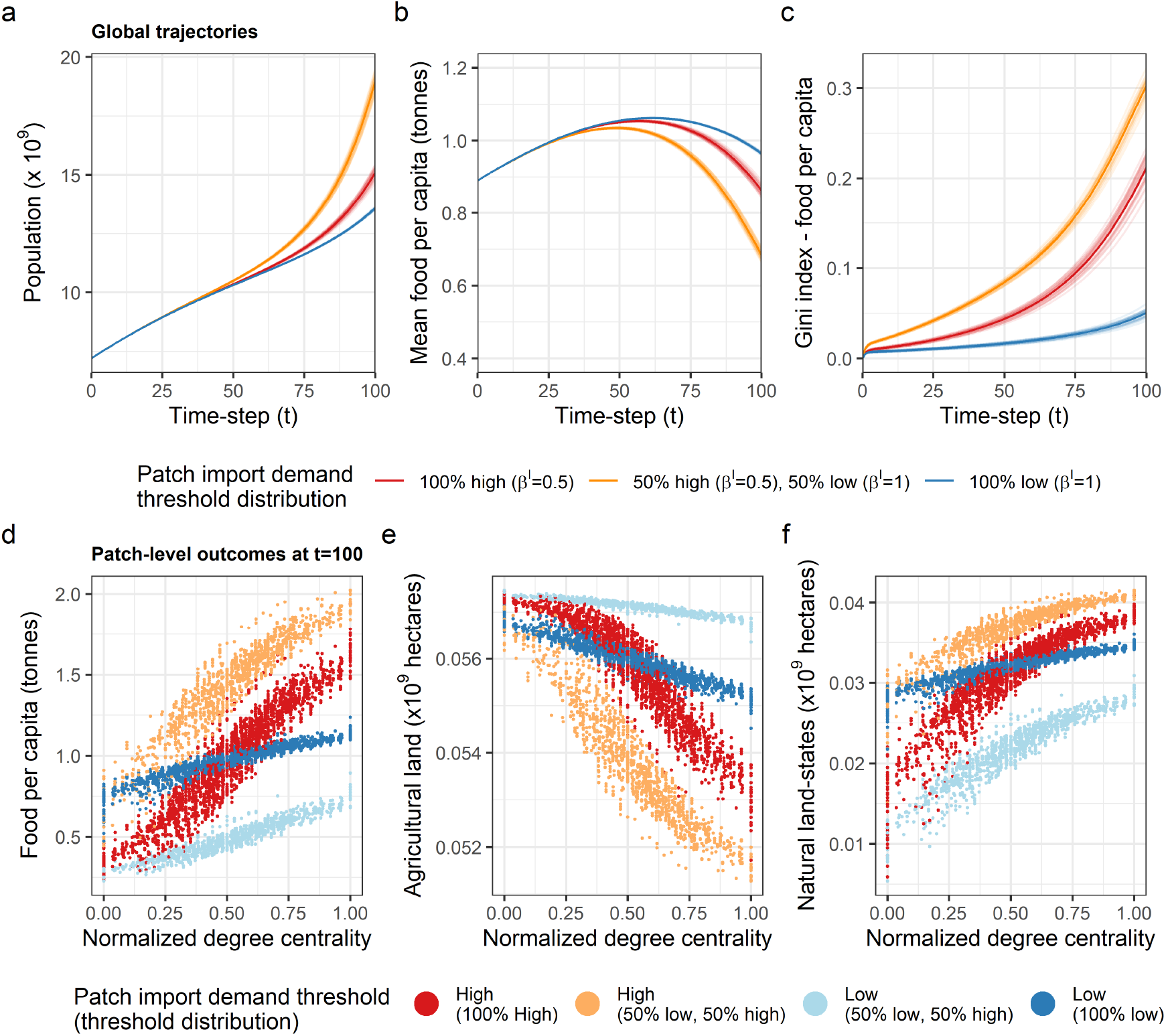
Patches that import food sparingly (i.e. have low import demand thresholds) lose out to patches that import heavily (i.e. have high import demand thresholds), exacerbating inequalities resulting from differences in network centrality. Figure panels show how patch-level import demand thresholds impact a) global population b) mean patch-level food per capita c) Gini index (food per capita), as well as how import demand thresholds and node centrality influence patch-level d) food per capita e) agricultural land area and f) natural land state area at *t* = 100. In a)-c) transparent lines indicate individual model realizations and solid lines indicate mean outcomes, in d)-f) points correspond to individual patches within model realizations. Model parameter settings (except for *γ* = 7.5, *β^A^* = 0.75, *β^I^*) appear in Supplementary information, Table S1.

We also consider how the import demand threshold interacts with network centrality to influence patch-level outcomes (Fig. 5d,e,f). Regardless of the parametrization, patches that are more central to the network (i.e. situated on nodes with high degree centrality values) obtain higher food per capita levels than their more peripheral counterparts, despite maintaining less agricultural land area, and retaining larger areas of natural land states. Centrality plays a larger role in determining food per capita and agricultural land area outcomes for patches that import heavily than for those that import sparingly (Fig. 5d,e). We see a similar effect on natural land state area when all patches have the same import demand threshold (Fig. 5f).

A scenario where all patches import sparingly results in the most preferable outcomes; mean levels of food per capita are highest, inequality is lowest, and all patches obtain food per capita similar to current levels (0.89 tonnes per capita [30]). Additionally, all patches retain relatively large areas of natural land states, whereas in other scenarios this is only possible for patches that are central to the network and/or import heavily.

To further explore the impact of heterogeneity, we vary the proportion of patches that import heavily. Inequality is highest when approximately 60 – 70% of patches import heavily (Supplementary information, Fig. S9). It becomes increasingly advantageous to import heavily as the proportion of patches that do so decreases. The fewer of them there are, the higher the food and non-agricultural land per capita these patches with high import thresholds can attain. Additionally, they retain larger areas of natural land states, in part due to their smaller agricultural land area. Patches that import sparingly also benefit as the proportion of patches importing heavily decreases, though to a lesser extent. Globally, natural land state area is maximised when there are no patches importing heavily. Similarly, the largest agricultural land area is obtained when there are very few patches importing heavily.

#### Demand responsiveness

Introducing heterogeneity in patch demand responsiveness (*γ*), where 50% of patches adjust their demand sharply in response to changes in food per capita (i.e. have high responsiveness) and 50% adjust demand gradually (i.e. have low responsiveness), results in a more unequal world (Fig. 6c) despite the fact that the global food supply is similar for all distributions of patch responsiveness (Supplementary information, Fig. S10).

**Figure 6.**
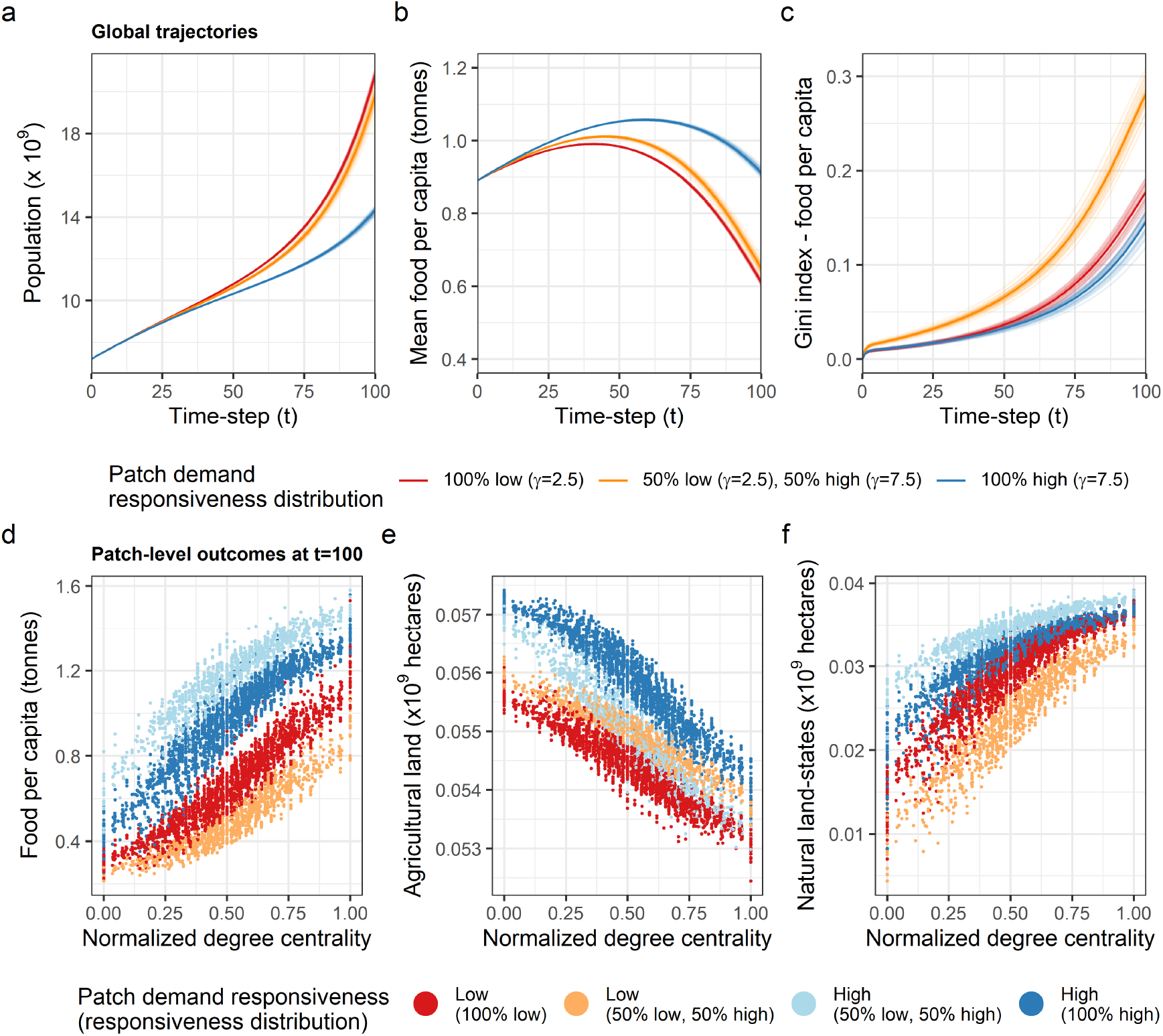
Patches that adjust their demand sharply in response to changes in food per capita (i.e. have high demand responsiveness) experience better outcomes than those employing gradual adjustments (i.e. those with low demand responsiveness). Figure panels show how patch-level demand responsiveness impacts a) global population b) mean patch-level food per capita c) Gini index (food per capita), as well as how demand responsiveness and node centrality influence patch-level d) food per capita e) agricultural land area and f) natural land state area at *t* = 100. In a)-c) transparent lines indicate individual model realizations, and solid lines indicate mean outcomes, in d)-f) points correspond to individual patches within model realizations. Model parameter settings (except for *β* = 0.75, *γ*) appear in Supplementary information, Table S1.

Initial increases in mean patch-level food per capita (Fig. 6b) are followed by a downturn, regardless of the distribution of responsiveness values. For all but the 100% high responsiveness case, this decline results in mean patch-level food per capita being lower at the end of the simulation than at initialization. Thus, though trajectories may initially appear promising, improvements in mean food per capita are unsustainable. This is most evident in the heterogeneously parametrized scenario where, on average, high responsiveness patches experience increasing levels of food per capita, low responsiveness patches experience decreases in food per capita in the latter stages of our simulations (Supplementary information, Fig. S11). The result is a large global population (Fig. 6a) in which the large food-poor populations of low responsiveness patches lay in stark contrast to the small food-rich populations of high responsiveness patches (Fig. 6d, Supplementary information, Fig. S11, Fig. S12).

Discrepancies in food per capita (Fig. 6d) caused by heterogeneous parametrization (i.e. differences in outcomes for 2 equally central patches with differing *γ*-values) are similar in magnitude to those resulting from differences in centrality (i.e. discrepancies in outcomes for the least and most central patches with the same *γ*-value). Centrality to the network facilitates the maintenance of lower levels of agricultural land area and large areas of natural land states (Fig. 6e,f). In a metapopulation where 50% of patches have high responsiveness and 50% have low responsiveness, there is little difference in the agricultural land area of 2 patches that are equally central to the network but display differing levels of responsiveness. However, the high responsiveness patch will tend to retain more natural land states (Fig. 6f).

The lowest level of inequality and highest mean food per capita are obtained when all patches are highly responsive to changes in food per capita. Here, the discrepancy in natural land state area between the least and most central patch is small. However, the most peripheral patches will still have substantially lower food per capita than their more central counterparts, with food per capita levels far below the current global average [30].

## Discussion

This study explored how the coupled dynamics of human population growth and food production systems are impacted by trade. We developed and analysed a simple metapopulation model expanding on previous work, e.g [24, 25], to incorporate land use dynamics and demographic transitions in human populations. We considered how the characteristics of trading entities (patches) and the trade network’s topology impact system outcomes. This discussion focuses on the linked ideas of equality and food security as facets of sustainability [13].

Metapopulations trading on larger networks experience less inequality. This result is likely due to the structure of small-world networks, where as network size increases, maximal path-lengths between nodes remain relatively low. This maintenance of short average path lengths facilitates increasingly efficient redistribution of resources with larger network sizes. Centrality to the network is key to attaining high food per capita. These results are consistent with those from a previous model [25].

A regular network structure, where all nodes are equally central, eliminates inequalities resulting from network topology. However, this structure may be impracticable for real-world networks. Countries specialize production, so certain goods will need to be imported even when food per capita is high [31]. There are costs (economic, environmental, etc.) associated with the transport of goods (e.g. creating and maintaining edges in the network) [16, 32]. Short average path lengths between nodes minimize the cost of obtaining these necessary imports while allowing for efficient redistribution of goods [33, 32, 34]. The structure of regular networks means that, unless they have a high edge density (with associated costs), they tend to have longer average path lengths than small-world networks of an equivalent size and density [35, 36]. If a country’s neighbours in a regularly structured trade network do not possess a good the country needs to import, the cost of doing so may become substantial. Small-world networks provide a compromise between efficient (and cost-effective) food distribution and the minimization of inequality.

When patch import demand thresholds are heterogeneous, patches that import sparingly (i.e. have low thresholds) are caught in a feedback loop between increased population growth and decreased food per capita. This phenomenon leads to outcomes where large populations and low food per capita persist in these patches. Patches that import sparingly may represent countries or regions for which it infeasible (financially, logistically, etc.) to import heavily, or those pursuing food security strategies emphasizing self-subsistence. The case of a metapopulation with heterogeneous demand responsiveness is similar, though the negative effects experienced by patches that gradually adjust their demand in response to changing food per capita (i.e. have low responsiveness) are less severe. Patches with low responsiveness may describe countries or regions where the agents (government, corporations, etc.) overseeing changes to agricultural land area and trade flows provide poor governance and/or are hampered by corruption.

Patches on the periphery of networks tend to bear more of the weight of agricultural production (i.e. maintain larger agricultural land areas). The uneven distribution of agricultural land area is particularly noticeable when patch import demand thresholds are heterogeneous; patches that import heavily have less agricultural land (and retain larger areas of natural land states) than their counterparts that import sparingly, despite maintaining substantially higher levels of food per capita. This result highlights the real-world phenomenon of a displacement or “leakage” effect, wherein countries increase imports to meet demand while protecting their natural land states. Displacement of agricultural land expansion to other countries results. If these countries have weaker protections for natural land states (e.g. displacement from developed to tropical countries) serious environmental impacts may be a consequence [3, 37, 16].

Strategies for improving equality and food security depend on the drivers of inequality. When all patches are identically parametrized, inequalities arise solely from differences in network centrality. Here, increasing parity in centrality by creating a more regular network structure improves equality and food security. When patch-level characteristics are heterogeneous, alterations to these characteristics provide an additional avenue for reducing inequality. This is particularly effective in the case of patch-level import demand thresholds, where the presence of patches that import heavily exacerbates inequalities created by differences in centrality. A shift towards all patches importing sparingly would improve food security and lead to a more uniform distribution of natural land states and agricultural land across patches. The latter effect would reduce the likelihood that ecosystems within peripheral patches will experience high stress due land conversion for agriculture or urban area.

Adjustments to patch-level characteristics may prove more expedient to implement, as these would not require large-scale network restructuring. However, such changes could require substantial sacrifices from individual patches. Network topology alterations may prove difficult due to the necessity of a coordinated effort by a large numbers of patches. Despite this, they may still be an appealing option due to their added benefits. A node’s (patch in our model) resilience to shocks and resource variability can be improved by increasing the number and/or diversity of its trade partnerships [38, 39, 25], and increasing the number of nodes in a network improves resilience at both the node and network level [25].

Future work could investigate how shocks to the network structure (resulting from armed conflict, trade wars, disease, etc.) [40] and/or climate change effects impact outcomes [25, 4, 7, 2]. Model realism may be enhanced by introducing more heterogeneity not only in patch-level characteristics but in the availability and quality of land suitable for agriculture. Allowing patches to shift their demand thresholds based on the relative payoffs of different threshold values could provide additional insight. Models for dynamic strategy switching in human populations are well established in the literature (e.g. [41, 42, 43]) and could be adapted. Other potential extensions include a dynamically evolving network [40], explicit consideration of a broader range of land uses [8], or temporally varying parameters [44].

Given the broad range of factors that must be accounted for when designing food systems that are resilient, equitable, and sustainable, modelling is a useful tool for identifying key areas for improvement. This work demonstrates the efficacy of simple differential equation models for exploring the dynamics of a human metapopulation linked through trade. Insights gained from simulating scenarios using such models will enhance our understanding of challenges for sustainable development.

## Methods

### Single patch model

We begin with a single-patch model of human population, agricultural land use, agricultural yield, and food supply, and we fit the model to empirical data to ensure that our single-patch dynamics are empirically plausible. Agricultural yield, *Y*, human population, *P*, agricultural land, *A*, and food supply, *F*, change according to

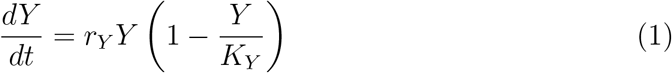

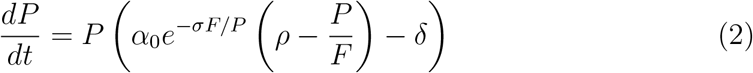

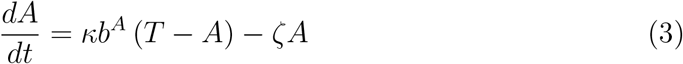

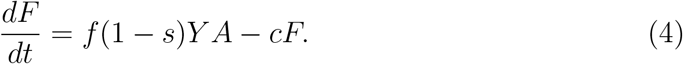

All parameter definitions are stated in Table S1. As in [8] we assume yield undergoes logistic growth. Food supply increases proportionally to agricultural yield and land area, and we assume that food supply is depleted over one year, on average (*c* = 1). Human population growth is density-dependent, with the birth rate changing dynamically with per capita food supply (details below). The total habitable land area, *T*, is the sum of agricultural land *A*, and non-agricultural land, which pools natural land states, urban area, etc.

Previous models have focused on resource-limited population growth, where increased access to resources raises the net population growth rate [24, 19, 25, 26]. The construction of our net human population growth rate function (Eq. 2) is intended to capture both resource-limited growth, and the occurrence of a demographic transition towards lower fertility rates when resources are sufficiently abundant. This phenomenon has been observed as countries become increasingly developed, leading to slowed population growth and in some cases a net population decrease [27, 28, 29]. The transition is likely driven by a variety of socio-economic factors; not only higher levels of wealth, but increases in the level of health, education, and other factors that are positively correlated with wealth [45, 28].

In order to incorporate this demographic transition into our model, which does not explicitly consider any measure of wealth, we utilize food per capita as a wealth proxy. We justify this use of a proxy by observing that more developed countries tend to have have higher per capita food supply, a consequence of food demand increasing with wealth – up to a point – as well as the higher levels of consumer-level food waste in developed countries [46, 47, 48, 49]. Thus, we are able to construct our net population growth function, assuming that in a resource-scarce regime food per capita is either the limiting resource or a proxy for wealth as the limiting resource, and in a resource-abundant regime it acts as a proxy for wealth. Combining these behaviours creates an overall curve, whereby the net growth rate increases to some maximum with increasing food per capita, prior to decreasing gradually at higher levels of food per capita. This curve resembles a Holling type-IV functional response [50, 51], and agrees with a previously proposed relationship between fertility and wages [52]. While there is some evidence to suggest that at very high levels of development (as measured using human development index) fertility will begin to increase, creating a “J-curve” [28], this result has been disputed [53].

Finally, we introduce a function, *b^A^*, that determines the response to changes in per capita food supply through adjustments to the rate of agricultural land expansion. This agricultural demand function is given by

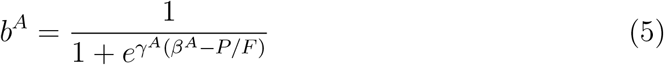

where *γ^A^* controls the steepness, and *β^A^* the midpoint of the sigmoid. In other words, *β* controls the demand threshold (smaller *β* indicates a higher food per capita must be achieved for demand to drop below 50%), and *γ* indicates the responsiveness to changes in food per capita (higher *γ* indicates larger change in demand for a given change in food per capita). With this function, as *P/F* increases (food per capita decreases), the rate of agricultural land expansion increases in response to increased demand.

Details of the fitting methodology carried out on the single patch model to obtain parameters for use in the metapopulation model can be found in the Supplementary methods. Single patch model trajectories using the parametrization resulting from this fit are presented in the Supplementary results.

### Metapopulation model

With all definitions as for the single patch model, we can formulate a metapopulation model of *N* patches as

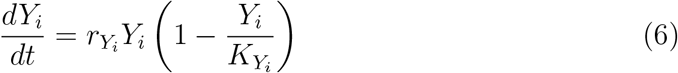

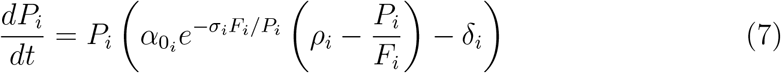

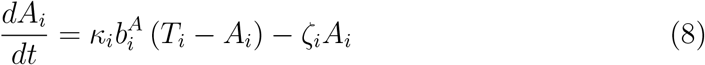

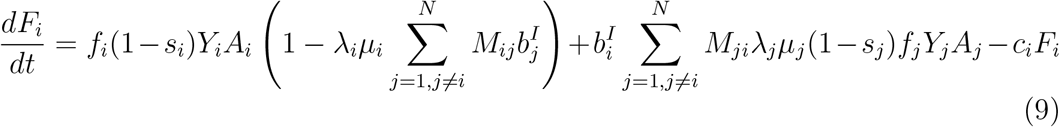

where *μ_i_* is the fraction of patch *i*’s production that is exported, with 1 – *μ_i_* consumed domestically. These patches may be thought of as countries, cities, or any other human metapopulation that exists within a trade network.

The (undirected, unweighted) trade network is represented as an *N*×*N* adjacency matrix *M*. We use undirected networks to describe trade, as the dynamic resource flows within the model will have some net direction without prior specification. Thus,

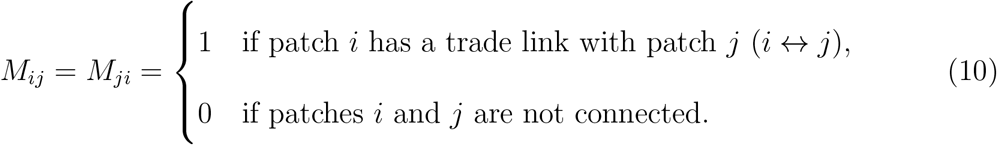

Empirical studies have shown that real-world trade networks are small-world [54, 34]. Thus, we use small-world networks as the basis for the trade networks describing metapopulation structure in our experiments. The “small-world” property indicates that the network displays high clustering (like a lattice) and a short average path length (like a random network) [35, 36]. We quantify the small-worldness of our networks using the small-world measure developed by Telesford et al. [36]. For the purposes of our analysis, we call a network “small-world” if the small-world measure (*ω*) returns *ω* ∈ [−0.5, 0.5] for that network [36]. For further information on the network science concepts and metrics used in our analysis please refer to the Supplementary methods.

We assume that, as with agricultural land expansion, the population of a patch will adjust import demand in response to changes in per capita food supply according to an import demand function given by

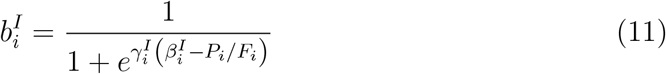

where 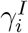 controls the steepness (i.e. the responsiveness), and 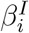 the midpoint (i.e. the demand threshold) of the import demand sigmoid for patch *i*. With this sigmoid function, as *P_i_/F_i_* increases (per capita food decreases), import demand increases.

We define a normalization factor, *λ_i_*, to ensure that a patch does not export more than a fraction *μ_i_* of its production. The value of *λ_i_* depends on external demand for patch *i*’s food, according to the relationship

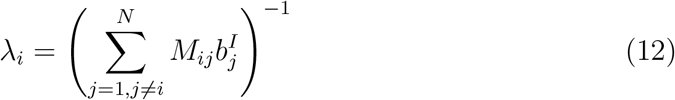

such that each trade partner is able to access an equal share of food, given their demand relative to the demand of all other connected partners.

Given these definitions for 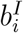 and *λ_i_* we can reduce Eq. 9 to

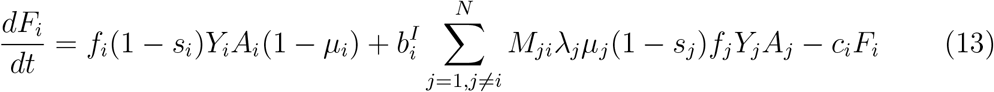

for simplicity.

All simulations for both the single patch and metapopulation models are run for 100 time-steps (years), with any results where the time-series is not explicitly referenced indicative of outcomes at *t* = 100. This time-frame is chosen as the assumptions that go into the construction and parametrization of models describing human behaviours are generally only valid over short periods, due to factors such as technological and societal change. For our model, these assumptions would include the constant parameter values for processes such as land conversion, as well as the use of a static trade network. The 100 time-step run was chosen as it has a similar length to the projections (to 2100) used by the United Nations and the Intergovernmental Panel on Climate Change to explore population growth, Shared Socioeconomic Pathways (SSPs) for climate change, and other phenomena [55, 1]. Considering this short time-span means that we are studying transient as opposed to equilibrium behaviour. However, given the rate at which change in human systems occurs, it is reasonable to assume that any equilibrium behaviour we could study by extending the time series would be invalidated as our model assumptions would no longer be reasonable.

Models are implemented in Matlab (Version 2016b), with equations solved numerically using ODE45 (adaptive 4^th^-5^th^ order Runge-Kutta method). In all experiments with the metapopulation model, we initialize the model by assuming global population, agricultural area, food supply, and available land are divided evenly amongst patches, in order to isolate the effects of trade. Using data from our single patch model (corresponding to the year 2013) under a low yield scenario as our *t* = 0 initialization we set *P_i_*(0) ≈ 7.20/*N* (×10^9^), *A_i_*(0) ≈ 5.00/*N* (×10^9^ hectares), *F_i_*(0) ≈ 6.41/*N* (×10^9^ tonnes) and *Y_i_*(0) ≈ 1.86 (tonnes/hectare). We initialize using model data instead of empirical data so that if the metapopulation model is reduced to a single patch model we will not see a discrepancy in results.

### Model analyses

To gain insight into how inter-patch trade interacts with patch-level dynamics to impact system outcomes, we perform several experiments manipulating different aspects of the trade mechanism and patch-level characteristics within the model.

#### Demand response

We first experiment with a simplifying case where the demand curves 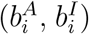 are identically parametrized 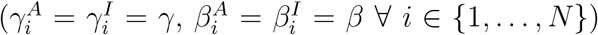 to explore the relationship between *β* and *γ*. Outcomes on the (*γ, β*) plane indicate how the system is impacted by the interplay between the demand threshold (controlled by *β*) and how responsive patches are to changes in food per capita (controlled by *γ*). This experiment is carried out under both “low” and “high” yield scenarios (see Supplementary methods for further details).

We also explore what happens when patch-level demand thresholds for domestic food (and thus agricultural land expansion) and imported food differ. This is implemented by setting 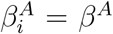 and 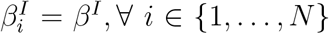 but *β^A^* ≠ *β^I^*. We assume that patches will have identical responsiveness to changes in food per capita 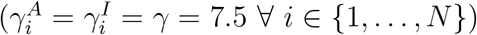 for both agricultural expansion and imports. At extremely low levels of food per capita 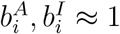 (scarcity necessitating a strategy of obtaining as much food as possible, regardless of source), and at extremely high levels of food per capita 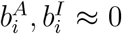 (as abundance removes the need to obtain additional food). At intermediate levels of food per capita, demand for agricultural land expansion within a patch will differ from the demand for food imports from other patches when 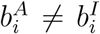. Here, a patch will keep importing food after it has stopped expanding agricultural land if *β_A_* > *β_I_*, or keep expanding its agricultural land after it has stopped importing food if *β_I_* > *β^A^*. This experiment is carried out under a “low” yield scenario, as are all subsequent experiments.

Both experiments are carried out on networks where the number of nodes (*N*) is either 10 or 100, to demonstrate results at multiple scales. We do not prescribe a set number of edges across all networks of the same size, thus allowing for a range of network densities. Instead we generate small-world networks with *N* nodes and a randomly selected neighbourhood size and rewiring probability. Once networks have been generated, 500 realizations of the model are generated for each of *N* = 10, *N* = 100, randomly selecting a network and (*γ, β*) or (*β^I^, β^A^*) pair.

#### Network structure

We explore how running simulations on networks with different structures (i.e. different levels of small-worldness) impacts outcomes both at the global and patch level. This re-structuring is done by rewiring regular networks using the Watts-Strogatz algorithm, to shift from more regular networks (when the rewiring probability, *p*, is low) through small-world networks (at intermediate rewiring probabilities) to more random networks (at high rewiring probabilities). For each rewiring probability, 25 networks are generated. We experiment with 100 node networks with either 1700 or 2600 edges, giving us network densities of *d* = 0.343 and *d* = 0.525 respectively. These densities are chosen to match those for the undirected real-world international agri-food trade network from the years 1986 and 2016 (the first and final year available in FAO trade data) as closely as possible [30]. We consider 2 densities to explore how the number of edges interacts with the rewiring probability, as previous work has shown the number of edges in real-world agri-food trade networks is increasing over time [40, 14, 56, 57].

For higher density networks we do not achieve highly random networks even when *p* =1 (Supplementary information, Fig. S13). Lower density networks experience more drastic and larger increases in the small-world measure, with the small-world measure becoming positive (indicating a network that is more random than regular) when *p* > 0.1. In lower density networks a higher rewiring probability leads to lower values for average path length and average clustering coefficient, while in the higher density network the average path length remains constant throughout. Networks at both densities experience changes in their average clustering coefficient over a similar range of *p*-values.

We limit ourselves to a 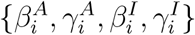 set that results in outcomes at *t* = 100 similar to those from our single patch model for the year 2100. This choice allows us to observe the effects of network structure when global outcomes align with plausible real-world trajectories. We choose 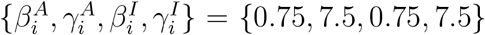, with 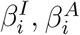 values taken from the region of (*β^I^, β^A^*) space where there is a moderate level of inequality (for our system and parametrization, where Gini index values rarely exceed 0.5) to better observe how network structures impact inequality (Fig. 3). For each combination of (*p, d*) there are 25 unique networks, and we run a single simulation on each.

#### Heterogeneous patch-level characteristics

We experiment with a heterogeneous parametrization of the model to ascertain how heterogeneity in patch-level characteristics impacts global and patch-level outcomes. These experiments are carried out on our 25 high-density networks (100 nodes, 2600 edges), with *p* = 0.5, so all results are taken across 25 simulations. This *p*-value is chosen as it is in the range of rewiring probabilities that result in small world networks (networks where the small-world measure is near 0, as shown in Supplementary information, Fig. S13). We assume these networks are a reasonable representation of real-world agri-food trade networks due to their small-worldness and large number of edges. We focus on degree centrality as our key network metric, as our analysis (see Supplementary results - Node centrality and patch-level outcomes) and previous work [25] suggest it is a consistent predictor of patch-level outcomes.

First, we examine the effect of heterogeneous import demand thresholds (*β^I^*-values). Here, 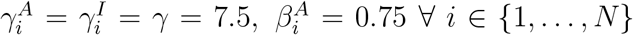, but 50% patches have a *β^I^* from the region of the (*β^I^*, *β^A^*) plane where outcomes between patches are highly unequal (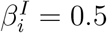, a high import demand threshold) while the remainder have 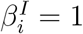 (a low import demand threshold) from a region with low inequality (Fig. 3mp). Patches with 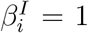 have much lower import demand (and thus import more sparingly) at a given level of food per capita than patches with 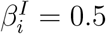 (except in the case where food per capita is either extremely high or extremely low). Patches are randomly assigned either the low or high 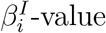, so there is no correlation between nodes based on patch import demand.

Second, we consider a simple case of heterogeneous responsiveness where 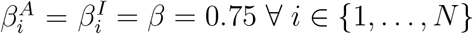, but 50% of patches have high responsiveness 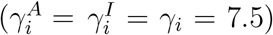 and 50% of patches have low responsiveness 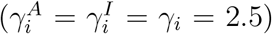. Highly responsive patches will adjust their demand for imports and agricultural land expansion sharply in response to changes in food per capita, while patches with low responsiveness will make more gradual adjustments. Patches are randomly assigned either the high or low *γ_i_*-value, so there is no correlation between nodes based on patch responsiveness.

#### Measures for describing outcomes

Across all experiments, we consider several measures, in addition to tracking the model variables, to enhance our understanding of food security and inequality in the system. Food per capita, already a driver of model dynamics, is also indicative of food availability, one of the 4 main pillars of food security. We also consider the amount of non-agricultural land per capita calculated as (total area – agricultural area)/person. We can interpret this metric as a measure of the per capita availability of natural land states as well as urban area for housing. Finally, we calculate remaining area of natural land states, based on the assumption that 0.06 ha per capita of land is required as urban area (the current global average [8, 58]), thus natural land state area is given by total area – agricultural area – 0.06*population.

Values for mean patch-level food and non-agricultural land per capita are weighted by patch-level population. Inequality between patches is described in the context of food per capita [59], and non-agricultural land per capita. We quantify these inequalities using the Gini index, which measures inequality between values within a distribution. It ranges between 0 and 1, with higher values correspond to greater inequality. Index values are calculated using the reldist package (Version 1.6-6) for R, with patch-level observations weighted by patch population size [60, 61]. We can think of high Gini index values for food per capita as indicative of inequalities in food security, while for non-agricultural land per capita these may indicate inequalities in access to housing, natural land states, etc. Units for our variables of interest and metrics are as follows; population (×10^9^ people), agricultural land (×10^9^ hectares), natural land states (×10^9^ hectares), food supply (×10^9^ tonnes), food per capita (tonnes/person), non-agricultural land per capita (hectares/person), Gini index (unitless).

## Data availability

Data sets required to run simulations are available in a GitHub repository (https://github.com/k3fair/MetapopulationTrade-model). Data sets generated from our analysis and simulations are available from the corresponding author upon reasonable request. All data used to parametrise the model are publicly available online [30, 1].

## Code availability

Code used for parameter fitting, simulations, analysis, and visualization is available in a GitHub repository (https://github.com/k3fair/MetapopulationTrade-model).

## Author information

### Contributions

K.R.F. conceived of the study, developed simulation code, performed all simulations and analysis, and wrote the first draft of the manuscript. All authors developed the model, designed the analysis, revised and commented on previous versions of the manuscript, and read and approved the final manuscript.

### Correspondence

Correspondence and material requests should be addressed to Kathyrn R. Fair.

