## Supplementary information for "Implications of trade network structure and population dynamics for food security and equality"

### Supplementary methods

#### Single patch model parametrization

The model is parametrized at the single patch (global) level using empirical data from the United Nations Food and Agriculture Organization (FAO). We collect data on global population ("Total population", 1961-2018, and projected "Total population", 2019-2100), food supply ("Food; the total amount of agricultural products available as food", 1961-2013), agricultural land ("Agricultural land; land used for cultivation of crops and animal husbandry. The total of areas under Cropland and Permanent meadows and pastures.", 1961-2017), total land area ("Land area; country area excluding area under inland waters and coastal waters", 1961-2017), uninhabitable land area (sum of "Terrestrial barren land" and "Permanent snow and glaciers" land cover areas from CCI-LI, 1992-2015), and food production ("Production quantity", excluding aquatic products, 1961-2013) [1]. We use a conservative estimate of the total land available for agriculture, taken as the total habitable land area (total area - uninhabitable area) in 2015, the most recent year for which all necessary data is available. While this undoubtedly leads to an overestimation of the amount of available land, the amount of land suitable for agriculture will vary widely depending on the production being considered (crops vs. livestock, type of crop, type of irrigation, etc.) [2, 3] so an accurate total across all production types is difficult to obtain.

To reduce the complexity of our fitting procedure, we independently fit the parameters from our net growth rate function (Eq. 2) as listed above the line in Table S1, to data on global net population growth rate and per capita food supply from 1962-2013. We assume $\delta = 0.0113$, the mean of the global crude death rate from 1950-2020, to focus our fitting on those parameters that do not have a direct biological interpretation [4, 1]. Through this assumption, we attribute all variation in the net population growth rate to per capita food supply driven changes to the fertility rate. We use a Levenberg-Marquardt type nonlinear least squares fitting algorithm implemented in R to obtain values for $\alpha_0$, $\rho$, and $\sigma$ (Table S1) [5, 6]. The resulting fit (Fig. S1) has a peak in net growth rate of 0.0343/yr when the annual food per capita is approximately 0.3673 tonnes. We note that the fitted value of $\rho$ is located at the upper bound of possible values ($\rho = 6.25$) however we feel that allowing for a lower value would create an unrealistic lower bound on the per capita food values for which net growth is positive, as our fit here is already fairly conservative (based on a comparison to the country-level data, as shown in Fig. S1).

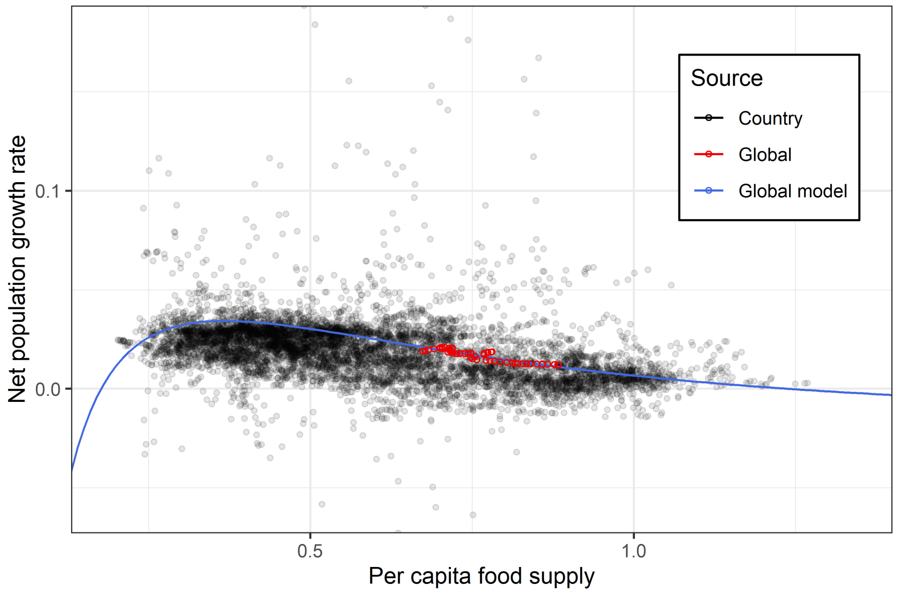


**Figure S1. Functional form developed for net population growth rate provides reasonable fit to empirical data.** Fitting is performed using global data on net population growth rate vs. per capita food for 1962-2013. Country-level data points shown for comparison.

We also obtain values for $r_y$ and $K_y$ to yield data prior to fitting parameters for the food supply and agricultural land (Eq. 3, Eq. 4), as the yield DE (Eq. 1) does not depend on the dynamics in the remainder of the system. For yield, we consider 2 scenarios; a "conservative" maximum yield scenario using the estimated maximum annual yield from [7] ($K_Y^C$=3.5), and a "high" yield scenario where we assume $K_Y^H = 2K_Y^C = 7$. We consider these contrasting scenarios as, given the roughly linear increase in yield from 1961-2013 and the potential for technological advances, the upper bound on yield may be higher than estimated by [7]. Alternatively, this "high" yield scenario can be thought of as a future where there is a global shift towards more plant-based diets, as livestock production requires a substantial amount of land and plant calories [8, 9, 10].

Once values for the parameters in the net growth rate function (Eq. 2), and yield (Eq. 1) have been determined, fitted values for the remaining parameters are obtained. The value of $\mu$ is fixed, as trade is not considered in the single patch model, meaning it cannot be included in our fitting exercise. We instead choose $\mu = 0.16$, the country-level average value for exports as a fraction of production in 2013, calculated using FAO data. All remaining parameters ($\kappa, \zeta, s, f, \beta^A, \gamma^A$) are fit under a low yield scenario. Fitting under a high yield scenario is also performed, where we assume that the fitted values of $\kappa, \zeta, s, f$ match those obtained under the low yield scenario and restrict our fitting to $\beta^A, \gamma^A$. All fitting (excluding fitting for the net growth rate function) is performed using the Matlab function lsqcurvefit applied to the single patch model. This function is a nonlinear-least squares curve fitting method utilizing the trust-region-reflective algorithm. [1].

**Table S1. Baseline parameters for metapopulation model.** Parameters related to human population growth (listed above line) fit independently of remainder of model fitting. Units: 1/year ($\delta$, $r_Y$, $\kappa$, $\zeta$, $f$, $c$), tonnes/(person $*$ year) ($\alpha_0$), tonnes/hectare ($K_Y$), tonnes/person ($\gamma^A$, $\gamma^I$), (tonnes/person)$^{-1}$ ($\sigma$, $\rho$, $\beta^A$, $\beta^I$), billions of hectares ($T$), unitless ($\mu$, $s$).

| Parameter | Description | Baseline value | Range | Source |
|---|---|---|---|---|
| $\delta$ | Base death rate | 0.0113 | N/A | [4] |
| $\alpha_0$ | Controls maximum net growth rate | 0.028 | $[0, 1]$ | fitting, [1] |
| $\sigma$ | Controls steepness of birth rate decrease with increasing food per capita | 2.1013 | $[1, 10]$ | fitting, [1] |
| $\rho$ | Controls food per capita where net growth becomes negative due to food scarcity | 6.25 | $[5, 6.25]$ | fitting, [1] |
| $K_Y$ | Yield carrying capacity (low, high yield) | 3.5, 7 | $[2, 7]$ | [7] |
| $r_Y$ | Yield growth rate (low, high yield) | 0.0301, 0.0245 | $[0, 1]$ | fitting, [1] |
| $\kappa$ | Agricultural land conversion rate | 0.0031 | $[0, 0.1]$ | fitting, [1] |
| $T$ | Total habitable land area | 9.6251 | N/A | [1] |
| $\zeta$ | Agricultural land abandonment rate | 0.001 | $[0.001, 0.1]$ | fitting, [11, 12, 13] |
| $f$ | Proportion of agri-food resources produced that are available as human food, annually | 0.75 | $[0.745, 0.755]$ | fitting, [14, 1] |
| $s$ | Proportion of food production unavailable due to pre-consumer food waste/loss | 0.0689 | $[0.02, 0.2]$ | fitting, [15] |
| $\mu$ | Proportion of food produced that is available for export | 0.16 | N/A | [1] |
| $c$ | Consumption rate, inclusive of consumer-level food waste/loss | 1 | N/A | N/A |
| $\beta^{A,I}$ | Control location of midpoint for agricultural land conversion and trade sigmoids resp. | N/A | $[0, 10]$ | N/A |
| $\gamma^{A,I}$ | Control steepness of agricultural land conversion and trade sigmoids resp. | N/A | $[0, 10]$ | N/A |

#### Network science concepts for analyses

We provide a brief overview of network science concepts required for analysis of our model. Our trade network is made up of nodes (on which the patches of our metapopulation model are situated) connected by edges (representing trade links between patches). Nodes are "neighbours" if they are connected by an edge. Throughout the remainder of the paper, when we refer to patch-level characteristics we are indicating properties of the dynamic behaviour in patch $i$, resulting from the parametrization of Eq. 6-Eq. 9. When we describe node-level characteristics, we refer to the metrics describing the position of node $i$ (that patch $i$ is situated on) within the trade network. The metrics we will employ to characterise nodes and networks are listed in Table S2. We include several centrality metrics to explore which measures of node "importance" (centrality, by various definitions) in a network have clear relationships to patch-level outcomes [16].

Throughout our analysis we generate networks using the Watts-Strogatz algorithm. This algorithm takes a ring lattice – a regular network, where the number of nodes and edges is fixed – and randomly rewires edges with some probability $p$. As the rewiring probability, $p$, increases the networks generated become less regular and more random. If $p = 0$ we retain the lattice, if $p = 1$ we have a completely random network, and for rewiring probabilities in $(0, 1)$, we may obtain small-world networks. This algorithm accepts as input a number of nodes, the "neighbourhood" in which to connect each node (equivalent to the average degree centrality of the network/2 for our undirected network), and the rewiring probability, $p$ [17]. All networks are generated using an implementation of the Watts-Strogatz algorithm from the igraph package (Version 1.2.4.2) for R (Version 3.6.3) [18, 6]. Network analysis is performed using both igraph the Network Toolbox package (Version 1.4.0) [18, 19].

**Table S2. Metrics for describing network topology.** A subset of common metrics from the field of network science useful to our analyses.

| Metric | Description | Source |
|---|---|---|
| Degree centrality | Number of edges connected to a node. | [20, 21] |
| Betweenness centrality | Extent to which a node is located on paths between other nodes. | [16, 21, 22] |
| Closeness centrality | Average distance from a node to all other nodes within a network. | [16, 21] |
| Eigenvector centrality | Measures influence in a network by accounting both for number and "quality" of edges, being connected to nodes with high eigenvector centrality scores contributes more to a nodes' eigenvector centrality than being connected to nodes with low scores. | [16, 23] |
| Local clustering coefficient | Average probability that a node's neighbours are themselves connected to each other. | [16, 21] |
| Density ($d$) | Ratio of the number of edges in a network to the maximum possible number of edges. | [24] |
| Average path length | Average length of the shortest paths (along edges) between pairs of nodes in a network. | [20] |
| Small-world measure ($\omega$) | Quantifies the extent to which a network displays small-world properties by measuring its average path length against that of a comparable random network, and its average clustering coefficient against that of a comparable lattice. This metric has a range of $[-1, 1]$ where values near 0 are indicative of a small-world network, more negative values point to a more regular (lattice-like) network, and more positive values point to a more random network. We consider networks to be small-world if $\omega \in [-0.5, 0.5]$, as in Ref. [25]. | [25] |

### Supplementary results

#### Single patch model

Our work with the single patch (global) model focuses on determining parameter values which allow the model to obtain a good fit to the empirical data we retrieved from the FAO (fitting method and data described in Supplementary methods - Single patch model parametrization). Fitting was carried out using data from the years 1961-2013 (additionally 2014-2100 for human population), under a low yield scenario, with $t = 1961$ as our initial condition. In the "low-yield" scenario (Fig. S2), our parameter fitting exercise indicates that the projected increasing global population trajectory will be accompanied by an increase in agricultural land area (and thus an increase in food supply). Fitted values for $\gamma^A$ and $\beta^A$ in this scenario are obtained ($\gamma^A = 10, /\beta^a = 0$) though any combination of $(\gamma^A, \beta^A)$ that results in $b^A = 1 \; \forall \; t \; \in [2013, 2100]$ will have the same outcome for variable trajectories.

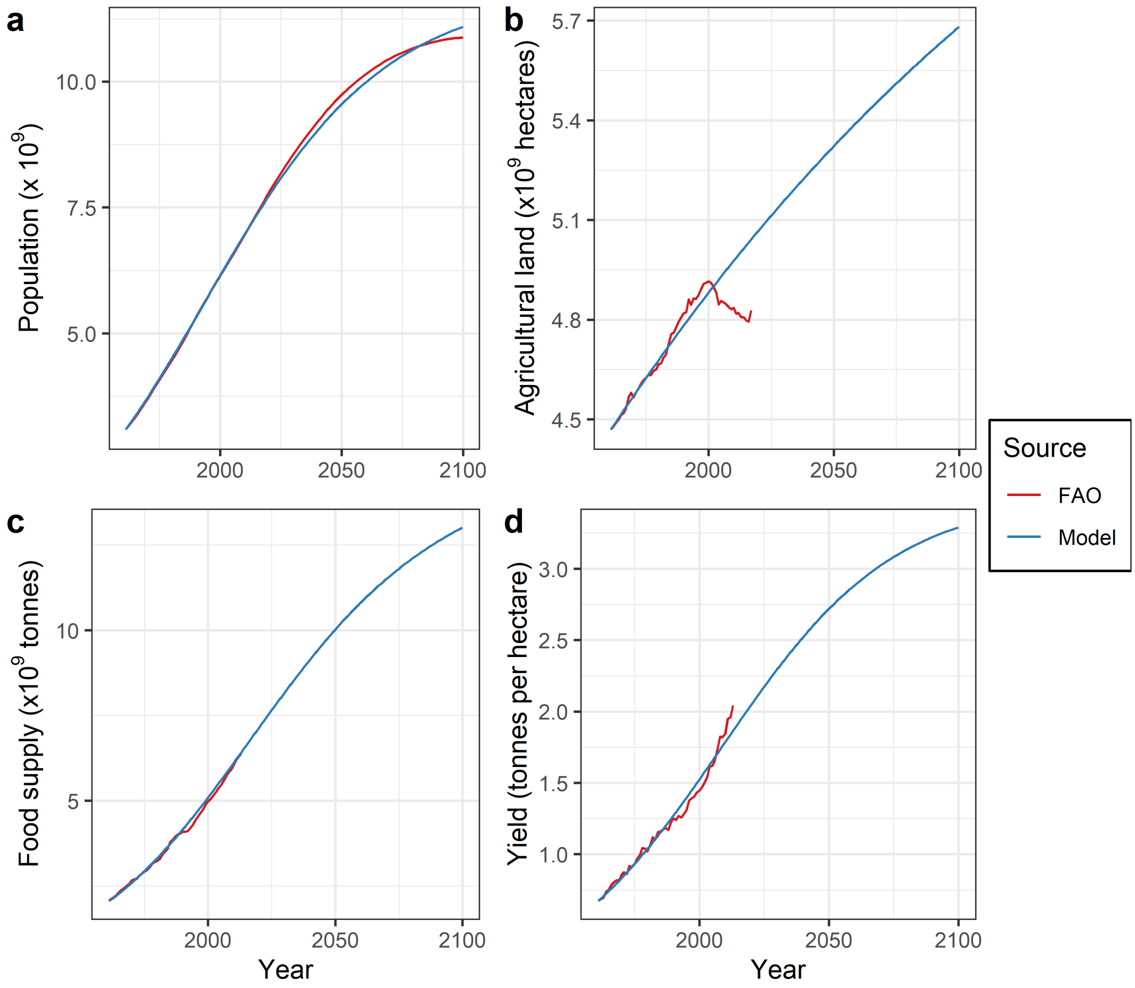


**Figure S2. Single patch model using low yield scenario provides reasonable fit to real-world trajectories.** Figure panels show trajectories for a) population b) agricultural land c) food supply and d) agricultural yield from the single patch model (low yield scenario) and empirical data.

In the "high-yield" scenario (Fig. S3), the higher value for maximum yield ($K_Y$) means that a larger food supply can be obtained, even with decreases in agricultural area past 2040. Additionally, this leads the population to experience a demographic transition to lower birth rates, causing a decrease in global population towards the end of the century. Here, a unique fit for $\beta^A$, $\gamma^A$ is obtained ($\gamma^A = 10, /\beta^a = 1.08$).

Though we obtain values for $\beta^A$ and $\gamma^A$ in these fits for the single patch model, we do not fix these values for our metapopulation model. This is motivated both by the lack of a unique fit in the "low-yield" scenario, and a desire to explore how $\beta^A$ and $\gamma^A$ interact with $\beta^I$ and $\gamma^I$. This exploration allows us to observe how the response curves for fulfilling food demand through expansion of agriculture ($b_i^A$) and imports ($b_i^I$) impact model outcomes.

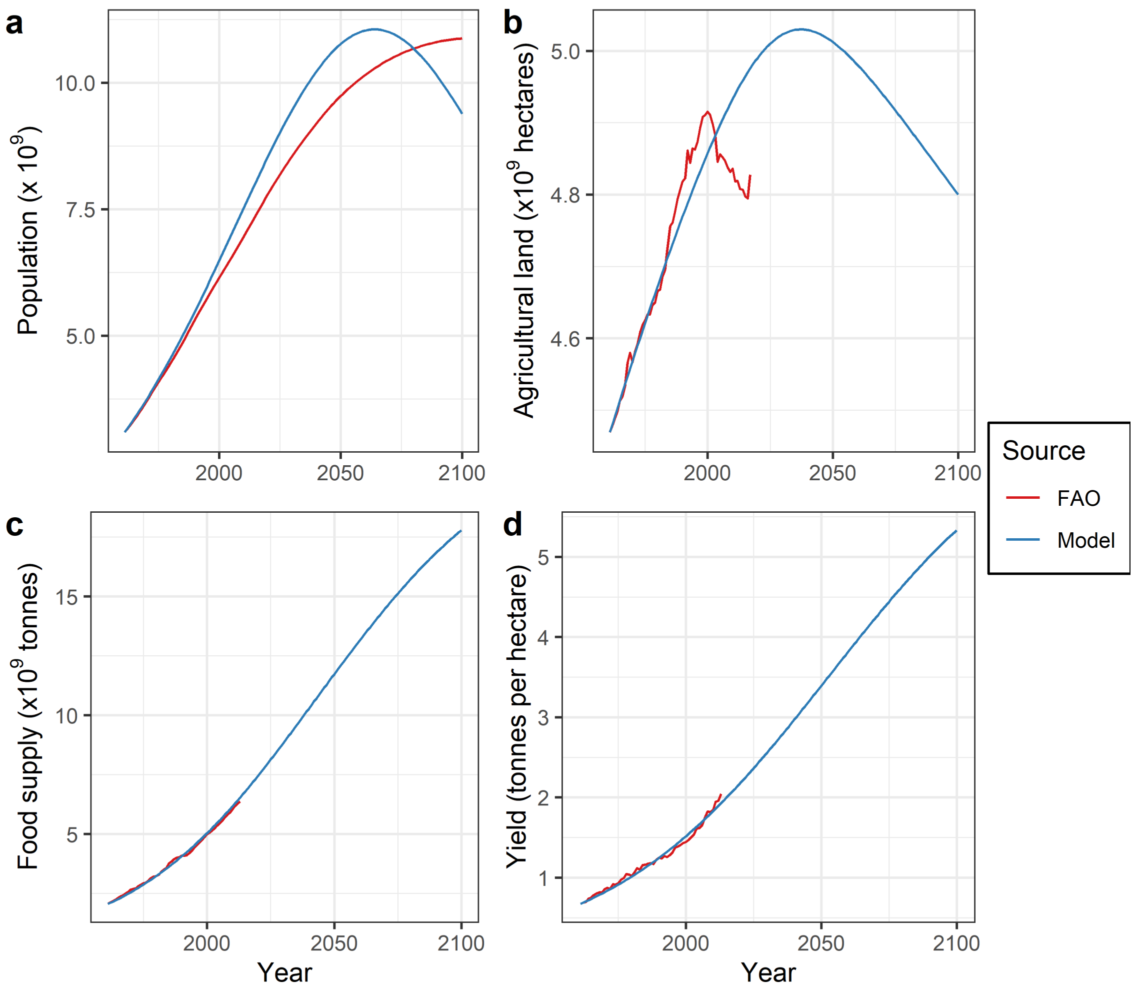


**Figure S3. High yield variation of single patch model alters trajectories to 2100.** Figure panels show trajectories for a) population b) agricultural land c) food supply and d) agricultural yield from single patch model (high yield scenario) and empirical data.

#### Node centrality and patch-level outcomes

Regardless of rewiring probability, more central patches (in terms of node degree, betweenness, closeness, or eigenvector centrality) will have higher food per capita at $t = 100$ (Fig. S4). This result is due to the fact that being situated more centrally in the network gives patches more access to food through imports. The effect on food per capita is reduced for networks generated using lower rewiring probabilities as these networks are more regular and differences in centrality between the most and least central node will be smaller.

It is unsurprising that all 4 centrality measures show similar results, as these metrics are highly correlated [26, 27]. However, unlike the centrality measures, local clustering coefficient does not appear to impact food per capita outcomes (Fig. S4e). Patches positioned on nodes with the same clustering coefficient experience a broad range of food per capita values, particularly when $p = 1$. Thus, being positioned on a node located in a highly clustered region of the network does not confer any advantage or disadvantage in terms of food per capita outcomes.

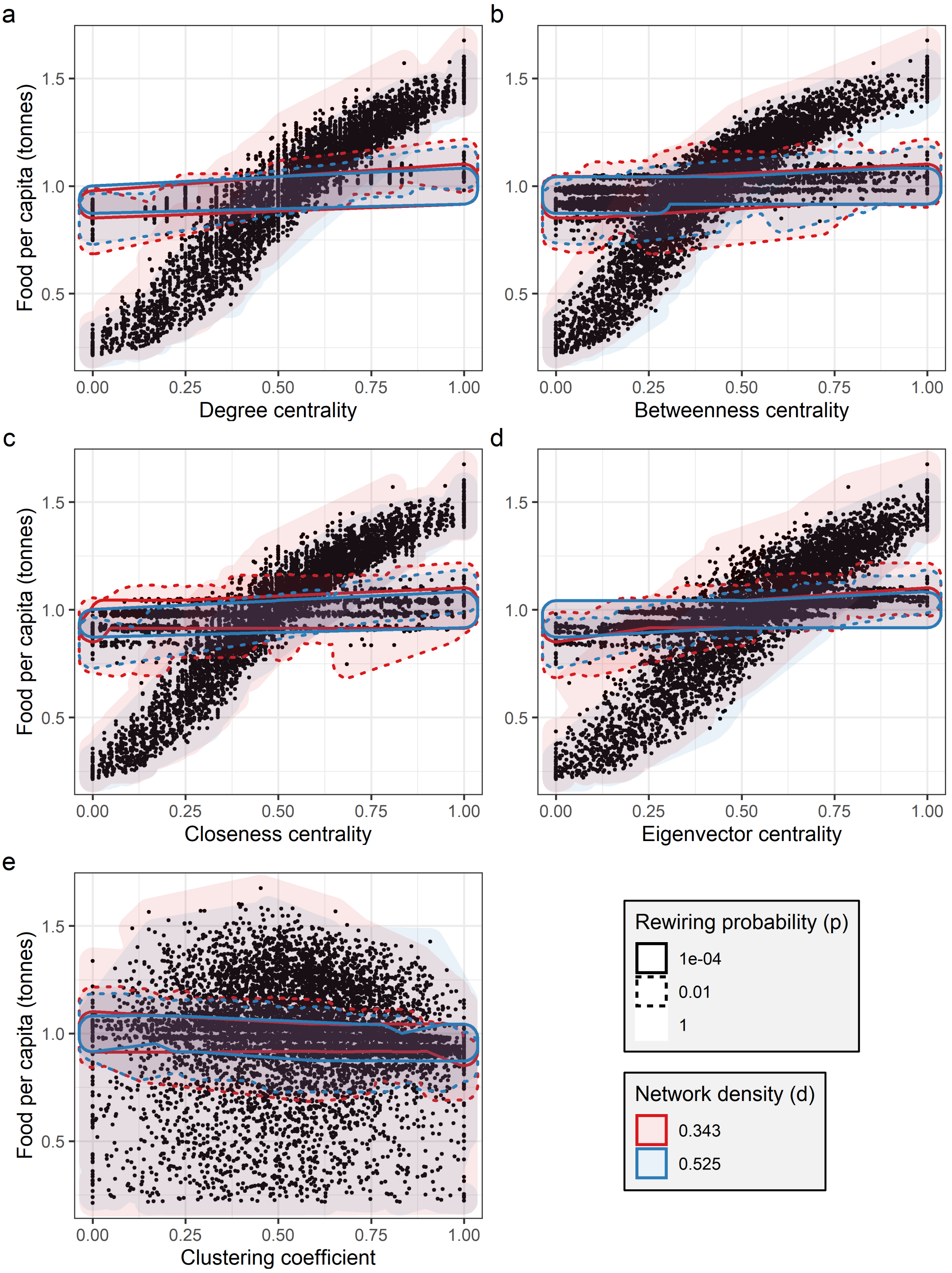


**Figure S4. Being more central to the trade network allows patches to obtain higher levels of food per capita.** Figure panels show the relationship between patch-level food per capita and a) degree centrality b) betweenness centrality c) closeness centrality d) eigenvector centrality e) clustering coefficient. All metrics are normalized to facilitate comparison across networks. Hulls are drawn around groups of points to indicate they correspond to nodes from networks with different densities and rewiring probabilities. Each point corresponds to a patch within a realization of the model. Model parameter settings (except for $\gamma = 7.5$, $\beta = 0.75$) appear in Table S1.

### Supplementary figures

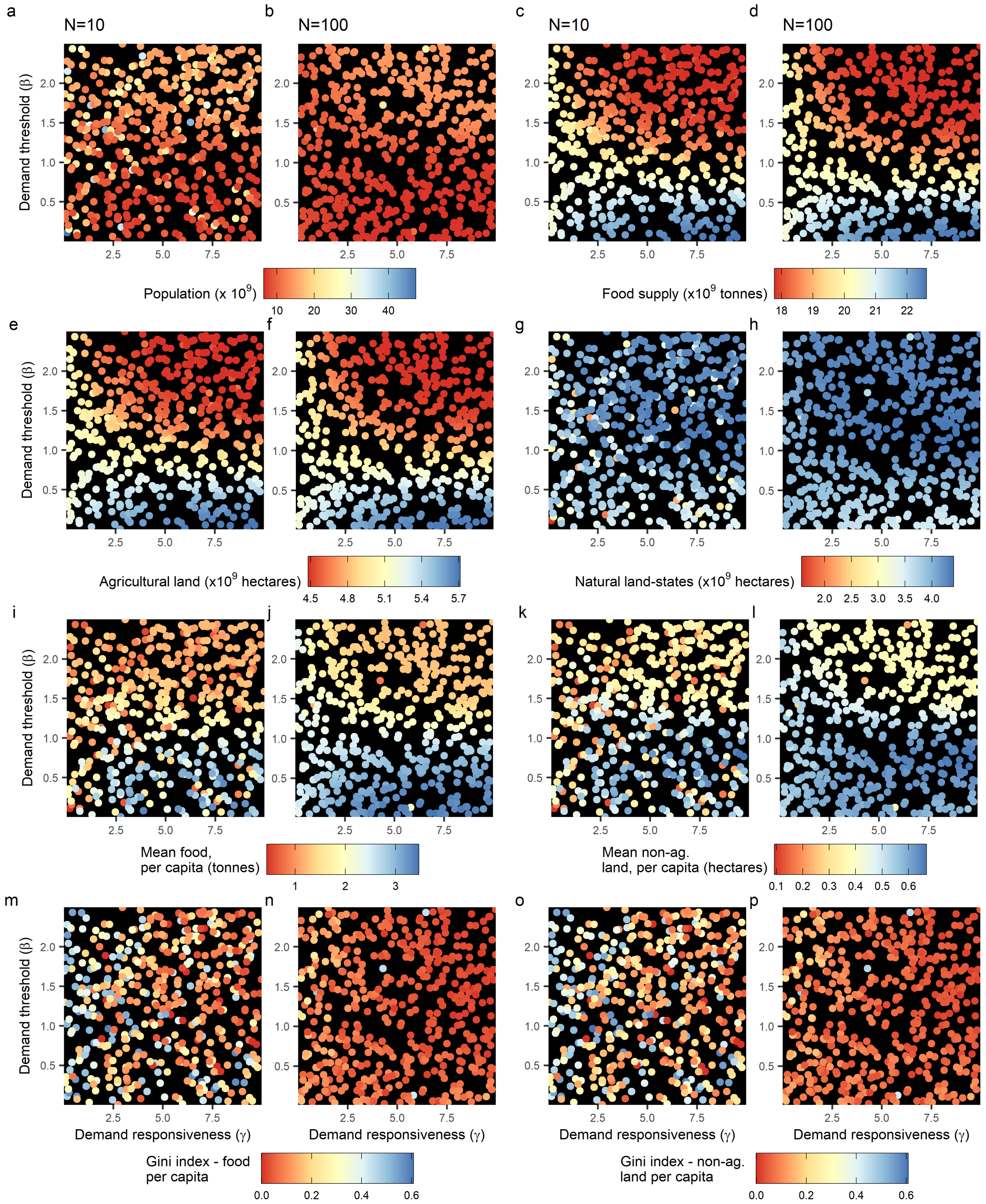


**Figure S5. Higher yields lead to a smaller global population, due to the transition to lower growth rates in food-rich populations.** Figure panels show how, under a high yield scenario, demand responsiveness ($\gamma$) and threshold ($\beta$) impact global a)-b) population c)-d) food supply, e)-f) agricultural land area, g)-h) natural land-state area, as well as mean patch-level per capita i)-j) food and k)-l) non-agricultural land, and Gini index, per capita m)-n) food and o)-p) non-agricultural land. rio. Each point indicates a realization of the model. Model parameter settings (except for $\beta, \gamma$) appear in Table S1.

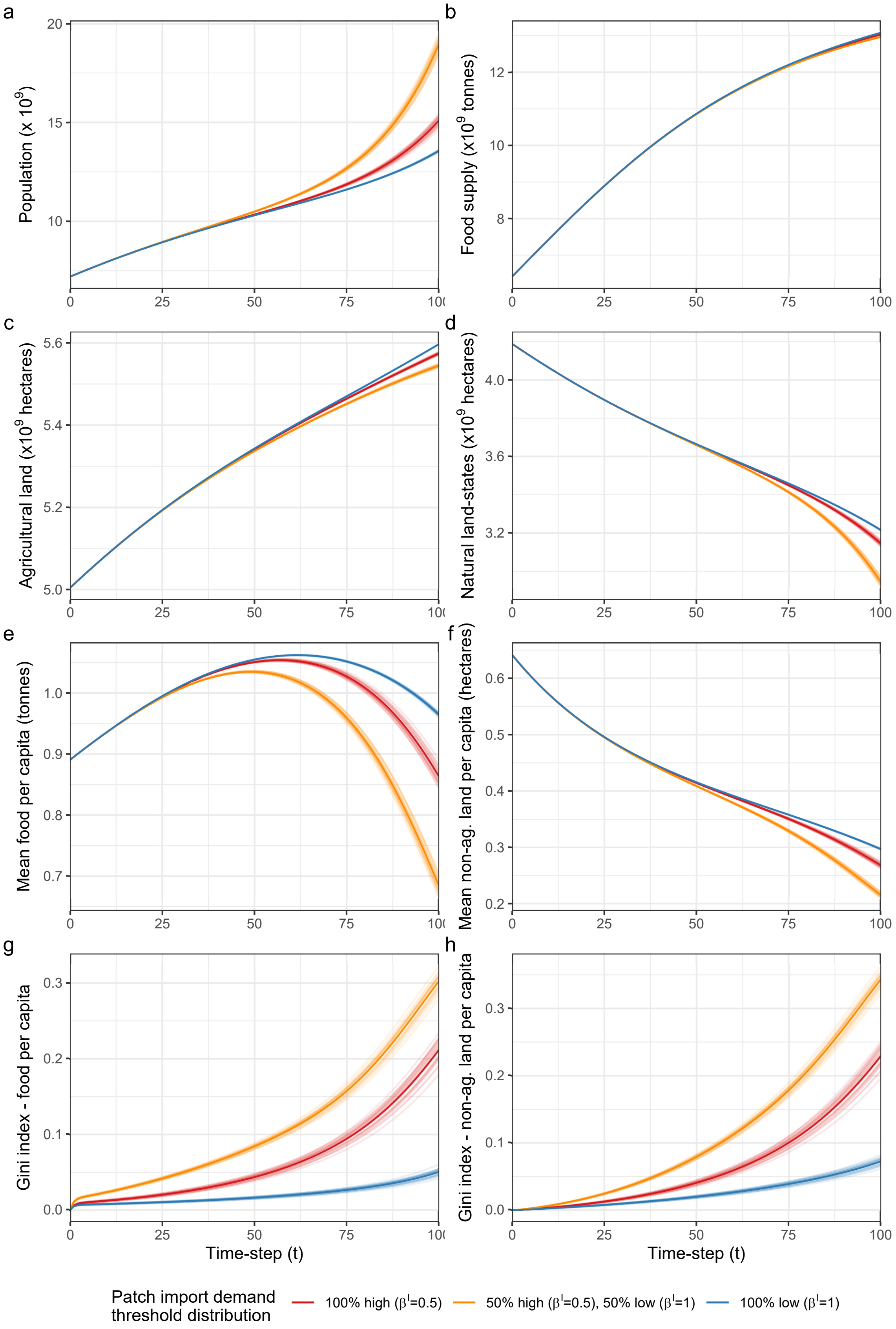


**Figure S6. Heterogeneity in patch import demand thresholds increases inequality.** Figure panels show the effect of heterogeneity in patch import demand thresholds on global a) population b) food supply c) agricultural land d) natural land-state area e) mean food per capita f) mean non-agricultural land per capita g) Gini index - food per capita h) Gini index - non-agricultural land per capita. Model parameter settings (except for $\gamma = 7.5$, $\beta^A = 0.75, \beta^I$) appear in Table S1.

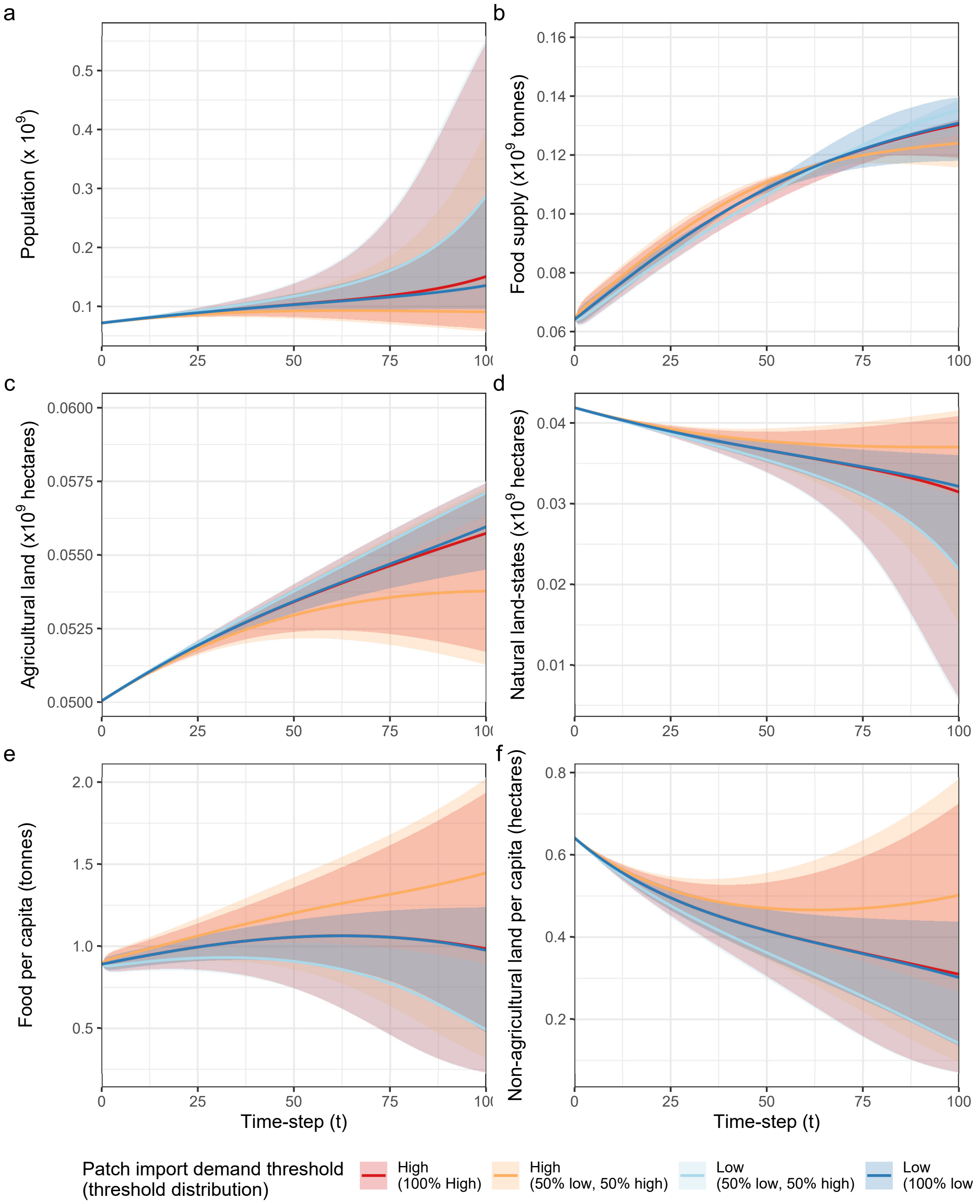


**Figure S7. Patches that import heavily (i.e. have high import demand thresholds) obtain substantially higher levels of food per capita than their counterparts that import sparingly (i.e. have low import demand thresholds).** Figure panels show the effect of heterogeneity in patch import demand thresholds on patch-level a) population b) food supply c) agricultural land d) natural land-state area e) food per capita f) non-agricultural land per capita. Lines indicate the mean outcome for patches by (patch import demand/import demand distribution) group, ribbons the range of possible outcomes (min/max values) across all patches in that group. Model parameter settings (except for $\gamma = 7.5$, $\beta^A = 0.75, \beta^I$) appear in Table S1.

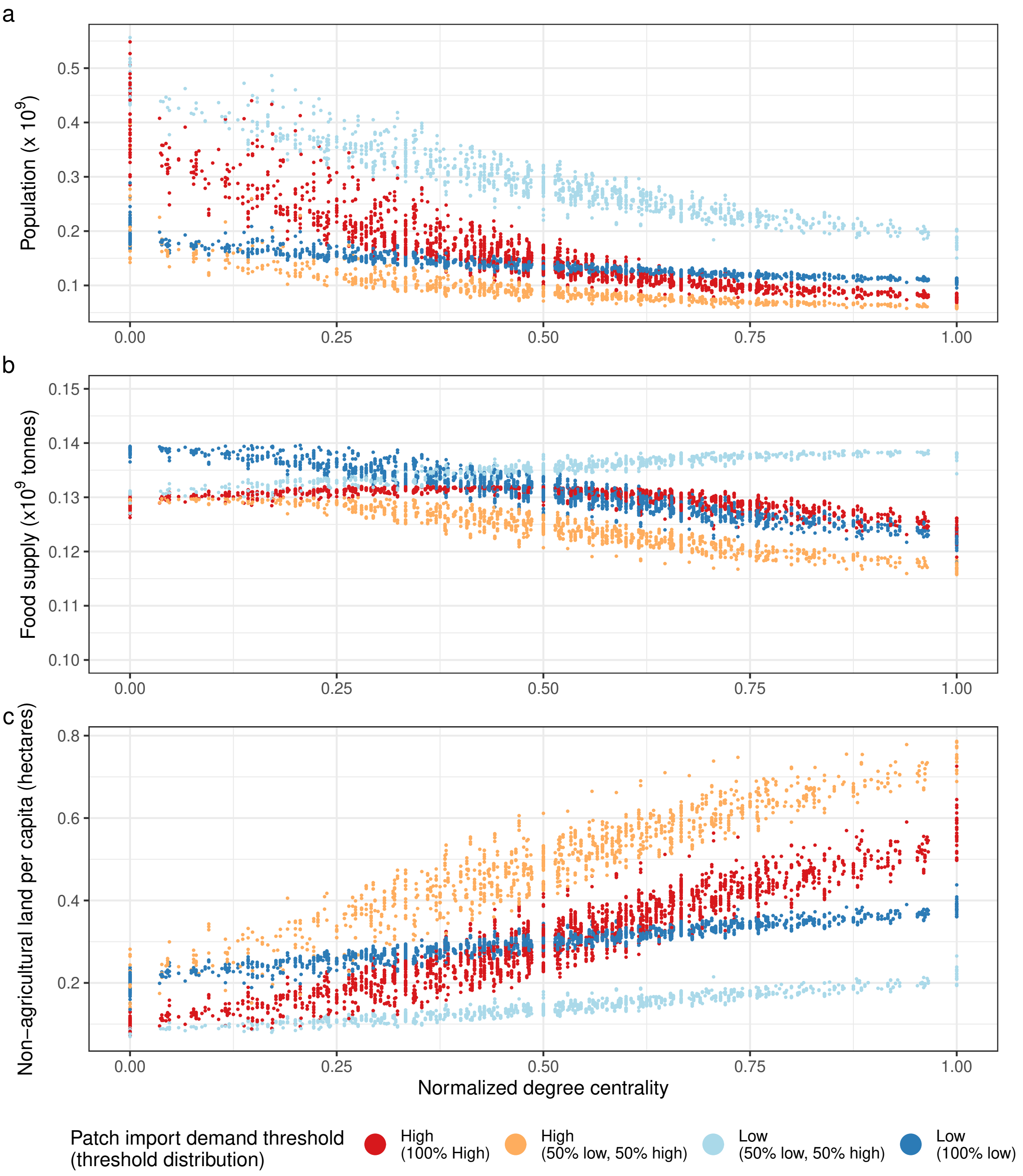


**Figure S8. Inequalities due to differences in node centrality are worsened when some patches import more heavily than others.** Figure panels show the effects of patch import demand thresholds and node centrality on patch-level a) population b) food supply and c) non-agricultural land per capita. Model parameter settings (except for $\gamma = 7.5$, $\beta^A = 0.75, \beta^I$) appear in Table S1.

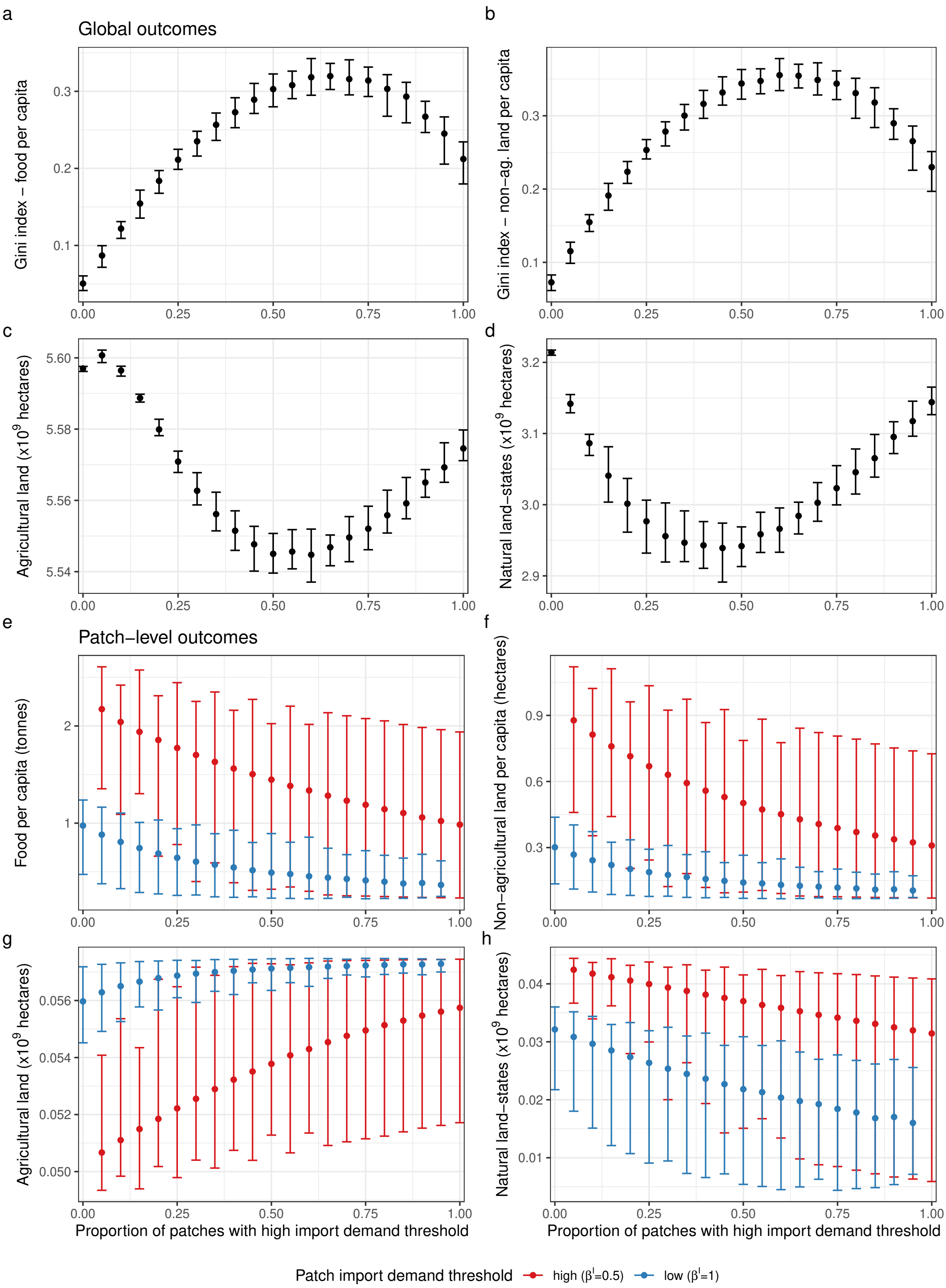


**Figure S9. Patches that import heavily (i.e. have high import demand thresholds) benefit most when they are rare.** Figure panels show how the proportion of patches with high import demand thresholds impacts a) Gini index - food per capita b) Gini index - non-agricultural land per capita c) agricultural land area d) natural land-state area, as well as the patch-level e) food per capita and f) non-agricultural land per capita g) agricultural land area and h) natural land-state area at $t = 100$. Bars indicate the range of outcomes (min/max values) and points indicate mean values across all realizations of the model. Model parameter settings (except for $\gamma = 7.5$, $\beta^A = 0.75, \beta^I$) appear in Table S1.

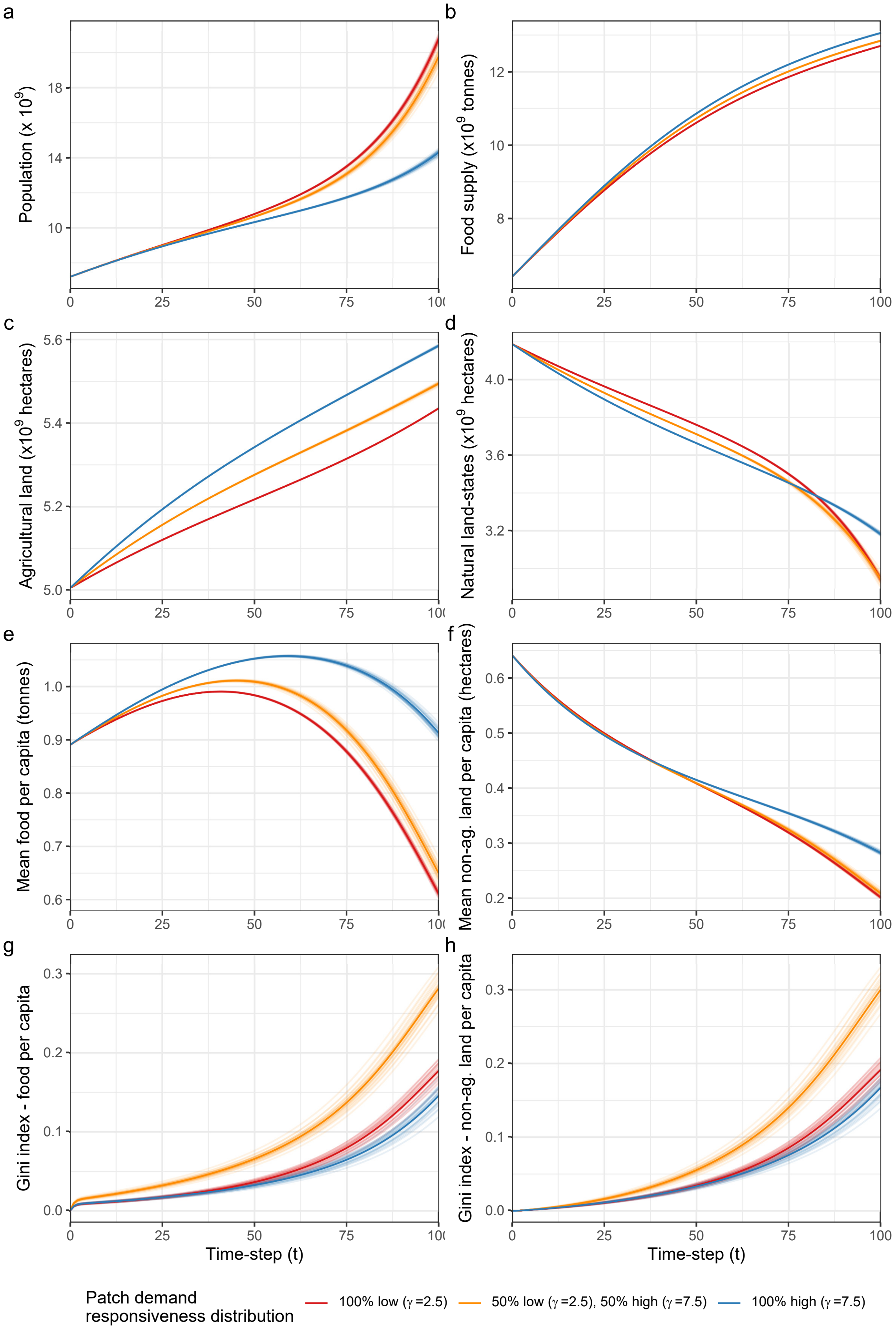


**Figure S10. Heterogeneity in how sharply patches adjust their demand in response to changes in food per capita heightens inequality.** Figure panels show the effect of heterogeneity in patch demand responsiveness on global a) population b) food supply c) agricultural land d) natural land-state area e) mean food per capita f) mean non-agricultural land per capita g) Gini index - food per capita h) Gini index - non-agricultural land per capita. Model parameter settings (except for $\beta = 0.75, \gamma$) appear in Table S1.

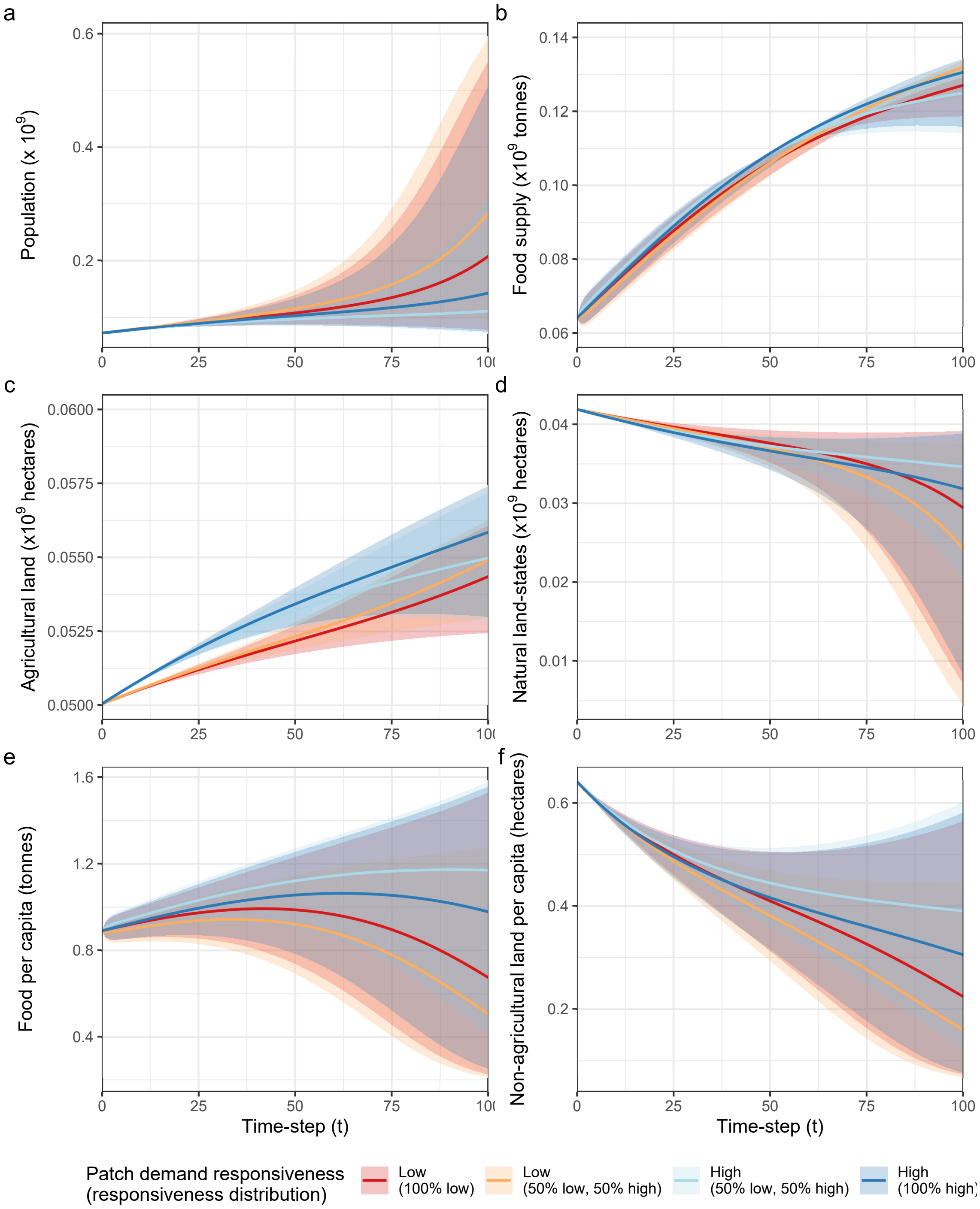


**Figure S11. Patches that adjust their demand sharply in responses to changes in food per capita (i.e. have high responsiveness) obtain better outcomes than their counterparts who respond more gradually (i.e. have low responsiveness).** Figure panels show the effect of heterogeneity in patch demand responsiveness on patch-level a) population b) food supply c) agricultural land d) natural land-state area e) food per capita f) non-agricultural land per capita. Lines indicate the mean outcome for patches by (patch responsiveness/responsiveness distribution) group, ribbons the range of possible outcomes (min/max values) across all patches in that group. Model parameter settings (except for $\beta = 0.75, \gamma$) appear in Table S1.

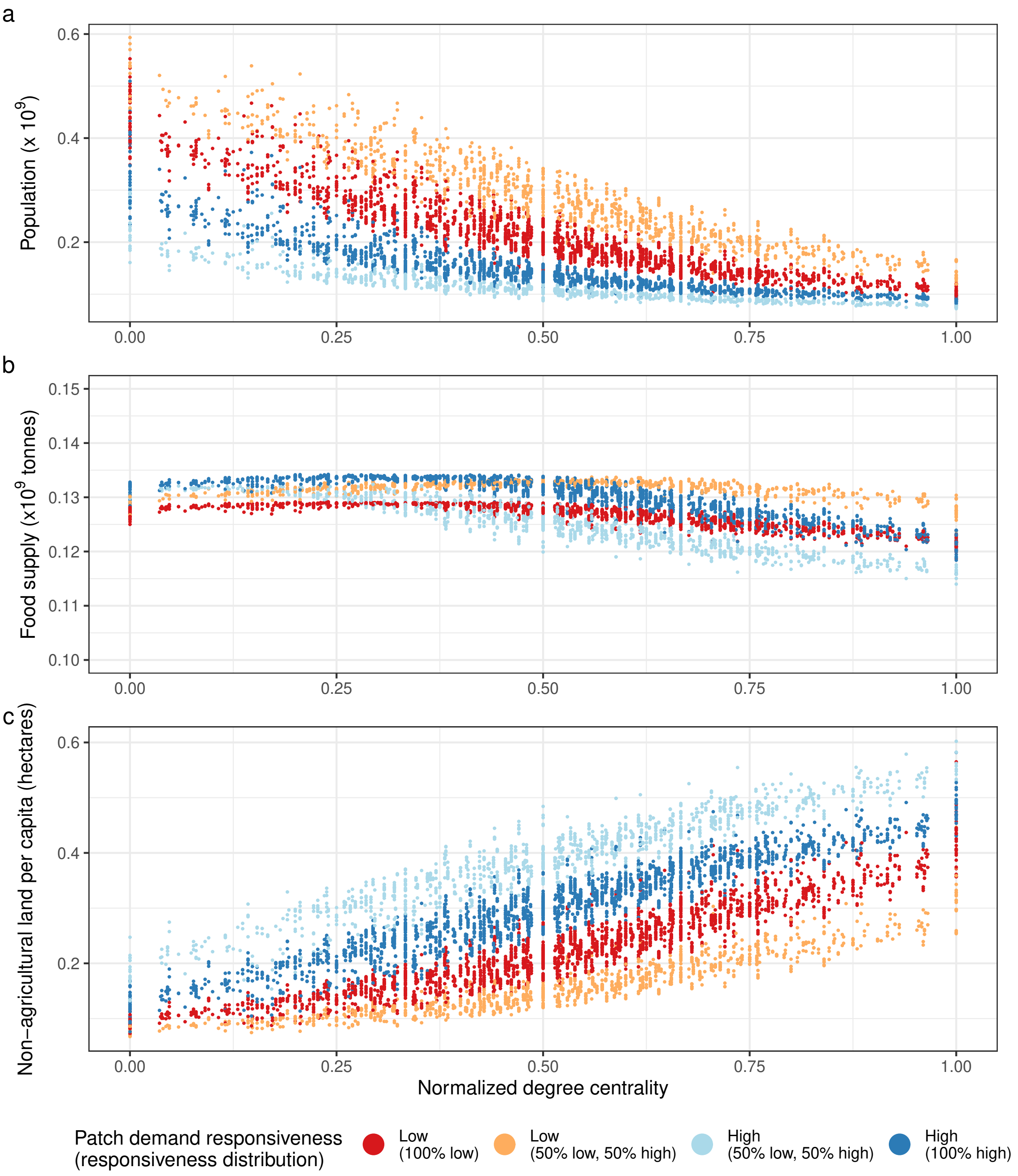


**Figure S12. Both differences in centrality and in how sharply patches adjust their demand in response to changes in food per capita contribute to inequality within the system.** Figure panels show the effects of patch demand responsiveness and node centrality on patch-level a) population b) food supply and c) non-agricultural land per capita. Model parameter settings (except for $\beta = 0.75, \gamma$) appear in Table S1.

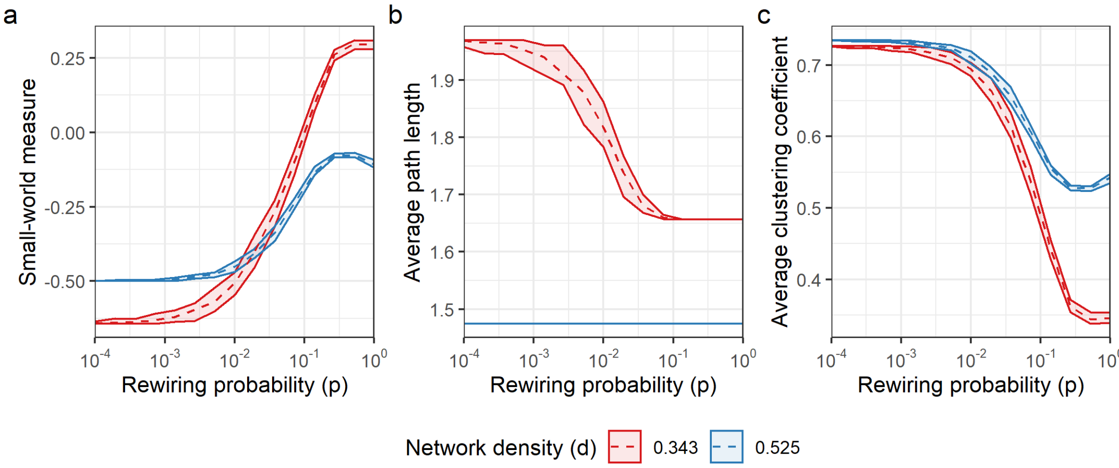


**Figure S13. Trade networks used for simulations exhibit the most small-worldness when $p \approx 0.5$.** Figure panels show the effect of the rewiring probability ($p$) on network a) small-world measure b) average path length c) average (local) clustering coefficient. Dashed lines indicate mean values for all networks with a given density and rewiring probability, while ribbons show the range of outcomes.

6. R Core Team. *R: A Language and Environment for Statistical Computing.* R Foundation for Statistical Computing, Vienna, Austria, 2020.

7. Chris Pagnutti, Chris T Bauch, and Madhur Anand. Outlook on a worldwide forest transition. *PLoS One*, 8(10):e75890, 2013.

8. David Tilman, Christian Balzer, Jason Hill, and Belinda L Befort. Global food demand and the sustainable intensification of agriculture. *PNAS*, 108(50):20260–20264, 2011.

9. Henning Steinfeld, Pierre Gerber, TD Wassenaar, Vincent Castel, Mauricio Rosales, Mauricio Rosales, and Cees de Haan. *Livestock's long shadow: environmental issues and options.* Food & Agriculture Org., 2006.

16. Mark E.J. Newman. *Networks: an introduction.* Number 134. Oxford University Press, 2010.

17. Duncan J Watts and Steven H Strogatz. Collective dynamics of 'small-world' networks. *Nature*, 393(6684):440, 1998.

18. Gabor Csardi and Tamas Nepusz. The igraph software package for complex network research. *InterJournal*, Complex Systems:1695, 2006.
